# Induction of transposable element expression is central to innate sensing

**DOI:** 10.1101/2021.09.10.457789

**Authors:** Derek C. Rookhuizen, Pierre-Emmanuel Bonte, Mengliang Ye, Thomas Hoyler, Matteo Gentili, Nina Burgdorf, Sylvère Durand, Fanny Aprahamian, Guido Kroemer, Nicolas Manel, Joshua J Waterfall, Richard Milne, Christel Goudot, Greg J. Towers, Sebastian Amigorena

## Abstract

Evidence indicates that transposable elements (TEs) stimulate innate sensing pathways in various pathologies but it is not clear whether they are sensed during normal physiological responses. Here we show that, during activation with an exogenous pathogen associated molecular pattern (PAMP), dendritic cells (DCs) epigenetically remodel heterochromatin at TEs by depleting the methyltransferase *Suv39h1* and reducing histone-3 lysine-9 trimethylation (H3K9me3). TLR4 signaling activates TE expression to enhance innate responses through the DNA sensor cGAS. Cytosolic cGAS-bound DNA comprised LINE1 TEs as the predominant endogenous ligands. Concordantly, LINE1 inhibition attenuated the type-I IFN response to LPS and rescued influenza virus infection. We propose that in healthy cells, exogenous PAMPs epigenetically activate self-derived PAMPs (LINE1) that engage cGAS to enhance responses. These data explain why pathogens employ redundant and broad innate immune countermeasures, to suppress activation of host PAMPs and illustrate a hitherto unappreciated role for host genome-derived PAMPs in response to pathogens.

## INTRODUCTION

The ability to distinguish self from non-self is a central principle of immunity. Invading pathogens must be recognized as non-self to trigger an adequate response while self-antigens must be tolerated to avoid autoimmunity. Innate detection of pathogens depends on the recognition of pathogen associated molecular patterns (PAMP) by pattern recognition receptors (PRR). PRRs include the Toll-like receptors (TLR) that sample extracellular and endosomal spaces for PAMPs (*1*) and various sensors that detect RNA and DNA (*2*). Upon PAMP detection, PRRs trigger signaling cascades that induce transcription factors to activate defensive gene expression. A key set of early defense genes include the type 1 interferons (IFN-I) that bind the ubiquitously expressed interferon receptor (IFNAR1/2) to activate JAK/STAT signaling and downstream transcription factors. The subsequent expression of up to 2000 interferon stimulated genes (ISGs) induces a defensive state in infected and neighboring uninfected cells through complex and incompletely understood mechanisms (*3*).

Recognition of foreign nucleic acid is a key step in sensing of pathogens, particularly viruses. Host nucleic acid sensors recognise nucleic acid in a non-sequence-specific way. Chemical differences in RNA structure such as phosphorylated 5’ ends, rather than the 5’ cap found in mammalian mRNA, or long double-stranded (ds) RNAs, distinguish pathogen RNAs that are detected by the PRRs RIG-I or MDA5, respectively (*4, 5*). Upon ligand binding, RIGI and MDA5 converge on the adaptor MAVS that activates TBK1 to phosphorylate IRF3 (Hornung et al., 2006; Li et al., 2009). In contrast, DNA sensing depends on DNA localization, rather than chemistry. DNA is not expected in the cytoplasm, and cytoplasmic DNA is sensed by the DNA sensor cyclic GMP-AMP synthase (cGAS) to activate IFN production (*6*). Upon binding to DNA, cGAS produces the cyclic dinucleotide 2’3’cGAMP (di-cyclic GMP-AMP monophosphate) that binds the adaptor protein STING and induces conformational changes leading to IRF3 activation(*7, 8*). Therefore, both RNA and DNA sensing culminate in IRF3-driven transcription of type-I IFN.

The fact that nucleic acid sensing is not sequence specific blurs the fundamental distinction between self and non-self. Indeed, an expanding and complex literature describes dysregulation of various types of host-derived retrotransposons leading to activation of either RNA or DNA sensors, production of type-I IFN and/or inflammation (*9–13*). Retrotransposons are transposable elements (TEs) that are broadly characterized by the presence (in endogenous retroviruses, ERVs) or absence (in LINE and SINE) of long terminal repeats (LTRs). TE expression can lead to the generation of dsRNA (*14*) and TEs encoding intact reverse transcriptase can produce dsDNA copies from their RNA genome for *de novo* integration into host genomes (*14–16*). Alternatively, RT-defective TEs, such as SINE and ETn, coopt the reverse transcriptase of other TEs to generate dsDNA(*17, 18*). Therefore, expression of TEs can generate nucleic acids that act as endogenous PAMPs and possibly drive deleterious immune responses. Given this potential to cause disease, it is striking that nucleic acid sensors have not evolved to avoid undesirable triggering by TE. Rather, it is assumed that in normal tissues TEs should be transcriptionally silenced and epigenetic histone modifications, including H3K9me3, are known to repress TE expression (*19*).

However, comparison of TE silencing between closely related apes revealed highly evolved control mechanisms suggesting that regulation of TE expression rather than complete silencing may be important in normal physiology (*20*). In this paper, we hypothesize that regulated expression of TEs is, in fact, part of a normal innate immune response. We tested this hypothesis in the context of the prototypic innate immune response to Gram negative bacterial lipopolysaccharide (LPS) and the response to infection with the single-stranded negative sense RNA virus influenza A (IAV). We show that stimulation of the anti-bacterial (*1*) and anti-viral receptor (*21*) TLR4 by LPS rapidly depletes H3K9me3 at LINE1 and SINEB1 loci by reducing expression of the histone methyltransferase *Suv39h1* and concomitantly inducing TE transcription through MYD88 and TRIF signaling. We find that the consequent LINE1 expression boosts the type-I IFN response to LPS in dendritic cells (DCs) by providing nucleic acid ligands for the cGAS/STING DNA sensing pathway.

Interrogation of cGAS-bound DNA in LPS stimulated cells, by immunoprecipitation and high-throughput sequencing (HTS), identified LINE1 as a principal cytosolic cGAS ligand. Furthermore, targeted genome-wide silencing of LINE1 elements using a CRISPR-dCas9-KRAB system confirmed that LINE1 act as endogenous PAMPs to activate cGAS and contribute to the consequences of LPS sensing. Crucially, single cell RNA sequencing (scRNA-seq) revealed that several distinct TLR pathways can activate the same set of TEs, suggesting a common mechanism among diverse innate stimuli. Indeed, inhibition of LINE1 expression, SUV39H1 overexpression, and treatment with reverse transcriptase inhibitors all rescued IAV infection of DCs, demonstrating the impact of derepressed TE expression on the host response to a pathogen encounter.

We propose that stimulation of cGAS via expression of reverse transcribing TEs contributes to the normal physiological response to pathogen-derived PAMPs, exemplified here by LPS and IAV infection. Our work establishes a new model in which TE expression is a central component of inflammatory innate immune responses. This is important because it suggests that pathogens must suppress recognition not just of their own PAMPs, but also that of TE nucleic acids derived from the host, perhaps explaining why so many pathogens encode apparently redundant innate immune countermeasures. Further, it defines a novel approach for investigation of diverse inflammatory and autoimmune pathologies and provides a new impetus for therapeutic innovation.

## RESULTS

### LPS induces *Suv39h1* regulated TE expression

In order to understand the role of TE expression during normal innate immune responses, we treated WT bone marrow-derived dendritic cells (BMDC) with the prototypical TLR4 ligand LPS and performed high throughput mRNA sequencing (mRNAseq). LPS triggered robust TE expression that manifested as waves of transcription—for example, cluster one at two to four hours, and cluster two at six to 18hrs (Figure 1A). TEs from the three retrotransposon superfamilies were all differentially expressed (DE) at the individual element level upon LPS treatment (Figure 1J). We hypothesized that distinct TE families might be uniquely targeted for expression by LPS stimulation. Thus, we compared the proportion of elements in different families before and after treatment using the hypergeometric test that revealed specific enrichment of LINE1 and different ERV families whereas SINE families did not significantly enrich (Figure S1A). Therefore, LPS selectively induces transcription of distinct TE families.

**Figure 1.**
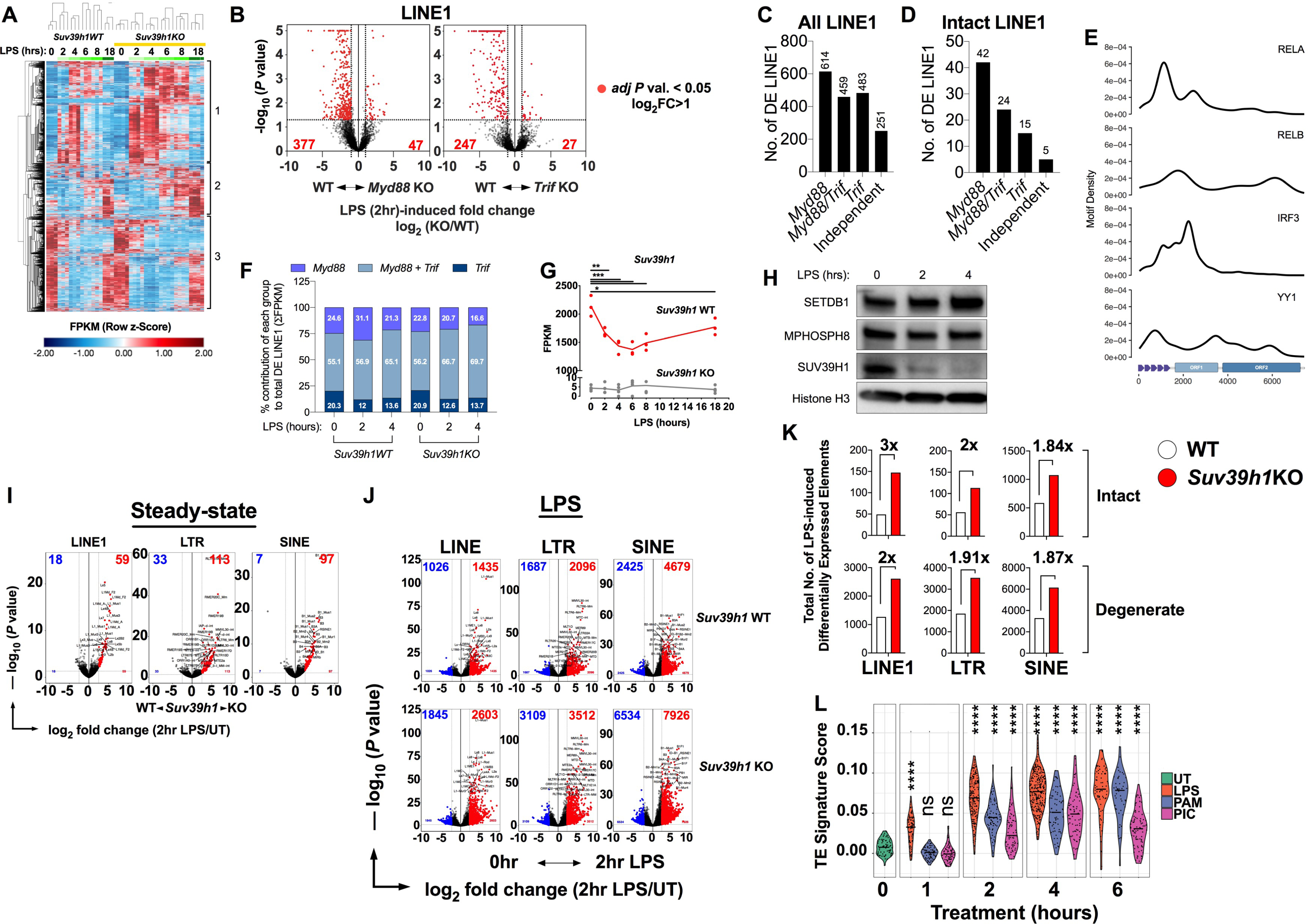
S*uv39h1* expression determines the magnitude of TE transcription during LPS challenge, see also Figure S1. (A) Heatmap showing Pearson Correlation-based hierarchical clustering of the most highly expressed TEs by normalized expression (FPKM) in WT (n=3) and *Suv39h1*KO (n=4) BMDCs. Relative values are represented by z-score. (B) Volcano plot exhibiting differentially expressed (DE) LINE1 elements in 2hr LPS-treated *Myd88^—/—^* (left) or *Trif^—/—^* (right) BMDCs compared with WT control cells. n=3 per genotype (*Myd88^+/+^*, *Trif^+/+^, Myd88^—/—^*, *Trif^—/—^*) per time point. Red dots signify significantly expressed elements, adjusted *P* value < 0.05. (C, D) Bar graphs representing the number of total (C) or intact (D) LINE1 elements dependent on the indicated signaling pathway for significant expression as determined by differential analysis of mRNAseq data such as that presented in panel B. (E) Density plots of IRF3, RELA, RELB, and YY1 transcription factor motif sequence (JASPAR database) enrichment along the sequence of full-length LINE1 elements. (F) Stacked bar plots showing the percentage of total DE LINE1 mRNA FPKM in *Suv39h1*WT and KO BMDCs at steady state and during LPS treatment that derives from each transcriptionally-defined group in panels C and D. (G) mRNA sequencing data showing that histone methyltransferase *Suv39h1* transcript levels are quickly depleted during LPS challenge in BMDCs. WT (n=3), KO (n=4). (H) Western blot of SETDB1, MPP8, SUV39H1, and Histone H3 in whole cell extracts from WT c57BL/6 BMDCs left untreated or stimulated with LPS for the indicated times. (I) Volcano plots displaying DE LINE1, LTR, and SINE element mRNA from sequencing data, comparing *Suv39h1*WT and *Suv39h1*KO BMDCs at steady-state. WT, n=3; KO, n=4. (J) Volcano plots displaying DE LINE1, LTR, and SINE element mRNA from sequencing data, comparing untreated (UT) to 2hr LPS-treated within WT or *Suv39h1*KO BMDCs. WT, n=3; KO, n=4. (K) Quantification of differentially expressed *intact* (top) or degenerate (bottom) TE classes in 2hr LPS-treated versus untreated within WT or *Suv39h1*KO BMDCs. (L) A TE signature score based on DE LINE1, LTR, and SINE in bulk mRNAseq data of WT BMDCs was projected onto a previously published scRNAseq dataset (Shalek et al., 2015) to compare TE expression induction by LPS, Pam_3_CSK_4_ (PAM), or poly(I:C) (PIC) treatment at the indicated time intervals. Significance cutoff for differential analyses in B-D and I-L was defined as an adjusted *P* value less < 0.05 and a log_2_ fold change ≥ 2. In C and D, the number of elements per group represents the sum of all significantly expressed LINE1 elements at 0, 1, 2, and 4hrs of LPS treatment when comparing WT to KO cells (*Myd88^+/+^*vs *Myd88^—/—^*, *Trif^+/+^*vs *Trif^—/—^*). In G, statistical significance was determined by one-way ANOVA and Bonferonni ad-hoc analysis comparing untreated to LPS-treated WT BMDCs. \**P*<0.05, ** *P* <0.001, \*\*\**P*<0.001, \*\*\*\**P*<0.0001.

Transcriptional control of TEs remains incompletely understood, particularly during normal physiology. TLR4 activates the canonical transcription factors *Nfkb* (*Rela* and *Relb*) and *Irf3* via the MYD88 and TRIF signaling pathways, respectively (*22–24*). The simplest hypothesis for LPS-induction of TE expression is that *Nfkb* and *Irf3* directly induce TE transcription. To test this, we performed differential expression analysis of LPS-induced TE transcripts in cells deficient for either *Myd88* or *Trif* (Figure 1B) and distinguished four groups of LPS-induced TE as follows—*Myd88* (*Myd88-*dependent)*, Trif* (*Trif-*dependent)*, Myd88-* and *Trif-*dependent (*Myd88/Trif-dependent*)*, Myd88-* and *Trif-*independent (independent) (Figure 1C, D, S1B). The *Myd88-, Myd88/Trif-*, and *Trif-*dependent groups contained the majority of LPS-induced, DE total LINE1 whereas the *Myd88-,* and *Myd88/Trif-*dependent groups selectively induced a majority of DE intact LINE1 (Figure 1D, see Materials and Methods for definition of intactness). Similar numbers of DE LTR elements were detected in the *Myd88, Myd88/Trif*, and *Trif* groups, and the *Trif* pathway selectively induced relatively more SINE elements (Figure S1B), distinguishing different modes of transcriptional regulation across the three retrotransposon classes. These observations reveal a previously unappreciated role for *Myd88/Nfkb* and *Trif/Irf3* in driving TE expression in addition to gene induction.

Our results indicate that autonomous TE expression is part of the normal transcriptional landscape downstream of classical LPS/TLR4 signaling. Concordantly, using LINE1 as an exemplar of TE, we found that RELA, RELB, and IRF3 binding motifs were enriched in the 5’ promoter region of intact elements (Figure 1E). As a positive control for our observations, YY1, a known LINE1 transcription factor in human (*25*) and mouse (*26*) also mapped to the LINE1 promoter region (Figure 1E). Altogether, our observations suggest that LPS directly regulates TE expression via canonical TLR4-induced transcription factors.

Although the *Myd88*-dependent group contained the most LINE1 elements (Figure 1C, D), it contributed a relativey intermediate amount to the total DE LINE1 mRNA, assessed by distribution of total LINE1 expression from mRNA-seq data (21-31%, Figure 1F). Meanwhile, the *Myd88/Trif-*dependent group contributed the largest proportion of total DE LINE1 mRNA (55-65%), whereas elements dependent on *Trif* alone contributed the least (12-20%, Figure 1F). Therefore, co-regulation by *Myd88* and *Trif* induced greater TE RNA expression, compared to either pathway alone. We conclude that both pathways contribute to induction of TE expression downstream of LPS detection by TLR4.

TEs are transcriptionally silenced by epigenetic modifications (*19*) such as H3K9me3 which is catalyzed by histone-lysine N-methyltransferases SUV39H1/2 and SETDB1, the latter cooperating with TRIM28. Previous work showed that SUV39H1 selectively trimethylated H3K9 at intact LTR and LINE1 loci in a mouse embryonic fibroblast cell line (*27*). Two studies have subsequently shown that, in cell lines, the HUSH complex also targets evolutionarily young TEs in collaboration with SETDB1 and TRIM28 (*28, 29*). Intriguingly, we found that LPS stimulation of BMDC rapidly and selectively depleted *Suv39h1* and *Setdb1* mRNA (Figure 1G, S1C).

*Trim28*, and HUSH complex members (*Morc2a, Mphosph8*) were also transiently decreased at the mRNA level. *Fam208a* (HUSH) and *Suv39h2* transcript levels were relatively stable in the early phase of the response but increased at later time points (Figure S1C). LPS treatment also induced several histone lysine demethylases belonging to the KDM family that target H3K9me3 (Figure S1C). Strikingly, *Suv39h1* was also lost at the protein level on LPS stimulation of BMDC whereas *Setdb1* and *Mpp8* were not (Figure 1H), indicating that additional mechanisms such as proteosomal targeting may impact *Suv39h1* expression. These results suggest that LPS stimulation rapidly remodels H3K9me3 heterochromatin with a key role for *Suv39h1*.

We therefore directly interrogated the effect of *Suv39h1* depletion on TE expression. In fact, mRNAseq revealed that knockout of *Suv39h1* in BMDCs (Figure 1G) significantly induced TE expression, even in the absence of LPS, as compared to WT cells (Figure 1I). Consistent with the notion that *Suv39h1* is a negative regulator of TE expression, LPS treatment of *Suv39h1*KO cells further enhanced expression of the same TEs induced by LPS in WT cells, with similar kinetics (Figure1A, J). For example, LPS induced 2603 differentially expressed LINE1 elements in *Suv39h1*KO compared to 1435 elements in WT (Figure 1J). We conclude that *Suv39h1* is a critical regulator of the magnitude of TE expression downstream of LPS stimulation.

Because evolutionarily younger (intact) elements have been implicated as PAMPs in diverse settings (*13, 30, 31*), we focused on the direct impact of *Suv39h1* expression on these elements specifically. Analyzing only intact elements reiterated that *Suv39h1* regulated expression of LINE1, LTR and SINE (Figure 1K). Specifically, *Suv39h1* KO had the greatest effect on DE of individual TE copies belonging to young LINE1 families (L1Md_F, L1Md_T, L1Md), to SINEB1, and to ERV1 and ERVK (Figure S1D-F), consistent with a previous study that used a mouse embryonic cell line (*27*). *Suv39h1* knock-out increased the number of DE intact LINE1 by 3 fold, upon LPS stimulation, compared with WT cells, whereas expression of truncated LINE1 elements was less impacted, approximately 2-fold (Figure 1K). Importantly, the expression of intact elements supports the model that the transcripts we are measuring represent autonomous TE transcription rather than transcriptional read-through of, for instance, intronic TEs or lncRNAs (*32, 33*).

Critically, the three groups of *Myd88/Trif*-regulated TE were represented in DE LINE1 elements in *Suv39h1*KO when compared to WT cells (Figure 1F), consistent with *Suv39h1* being a negative regulator of these elements. We hypothesize that LPS drives specific TE expression through simultaneous activation of *Myd88/Trif* to provide transcriptional activation via NFkb and IRF3 while concomitant loss of the negative regulator *Suv39h1* enhances TE transcriptional permissiveness. Thus, transcription factor specificity, and loss of H3K9me3 at TE loci due to SUV39H1 depletion, drive TE induction and determine the magnitude of expression.

We next used publicly available RNAseq data from ENCODE to ask whether LPS induces TE expression in human cells. Data from primary human monocyte-derived DCs demonstrated that, as for mouse BMDCs, untreated human cells expressed low levels of LINE, SINE, and LTR elements (Figure S1G, H) that were strongly increased by LPS treatment (Figure S1H). In particular, the largest effects were observed on LINE and SINE that peaked at two hours (Figure S1H). We saw that low levels of the relatively young, full length LINE1 L1PA elements and SINE transcripts were expressed at steady state but that LPS greatly boosted their transcription along with one copy of ERVK (LTR) (Figure S1I-K). Therefore, our observations of TE expression during inflammatory responses in mouse cells translate directly to humans with the difference that LTR expression may be relatively more restricted in humans.

Based on our analysis of transcriptional control of TEs by LPS, we hypothesized that additional innate sensors would induce TEs. Using differential expression of TEs in our bulk mRNA-seq datasets, we established a TE signature score, and used it to follow TE induction in a publicly available single-cell RNA-seq data set from BMDCs (*34*). This analysis revealed induction of the same TEs at single cell resolution as in bulk mRNAseq data following LPS treatment (Figure 1L). Stimulation of BMDCs with either a TLR2 (PAM_3_CSK_4_) or a TLR3 ligand (non-transfected dsRNA mimetic poly(I:C), PIC) that stimulate the *Myd88-* and *Trif-* pathways, respectively, induced robust TE expression (Figure 2J). Strikingly, these disparate PAMPs mimicking diverse pathogen stimuli, activated expression of the same TEs as defined by our LPS-based signature score, but with different magnitudes and kinetics of activation. Therefore, these data expand our observations with different protocols, and support our hypothesis that induction of TE expression is central to innate immune sensing. Altogether, these data suggest that physiological depletion of *Suv39h1* following LPS detection, selectively permits expression of TEs.

**Figure 2.**
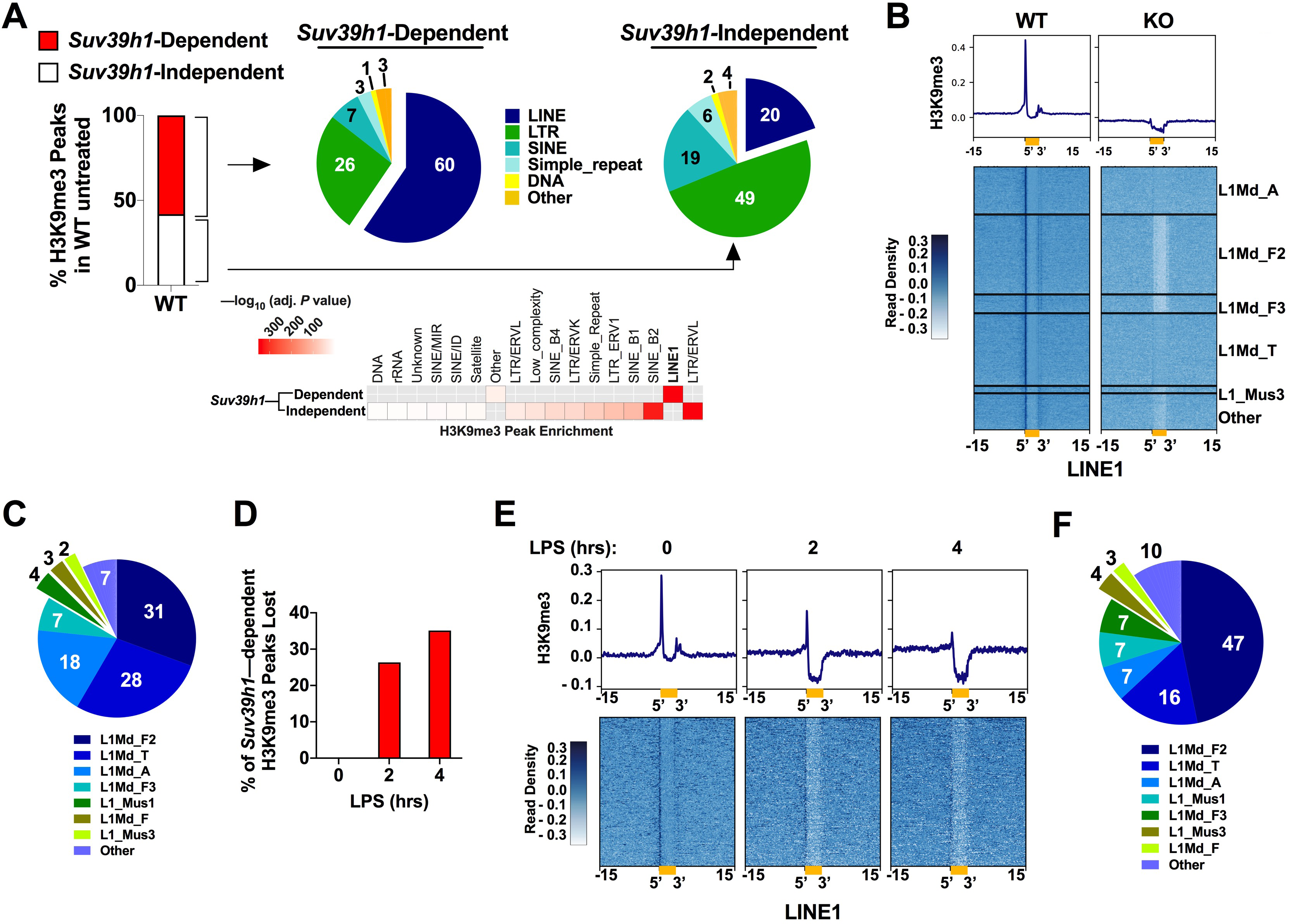
LPS induces rapid depletion of H3K9me3 at transposable elements in dendritic cells, see also Figure S2. (A) Left, stacked bar chart summarizing proportion of *Suv39h1*-dependent (red) and –independent (white) H3K9me3 peaks in unstimulated WT BMDCs from H3K9me3 ChIP-seq data. Right, pie chart exhibiting percent enrichment (% indicated by emboldened numbers) of specific DNA species in *Suv39h1*-dependent (middle) and –independent (right) H3K9me3 peaks. Below, heatmap summarizing statistically significant enrichment of DNA elements in *Suv39h1*-dependent versus *Suv39h1*-independent H3K9me3 peaks. (B) H3K9me3 ChIP-seq reads aligned by the 5’ end of intact *Suv39h1*-dependent LINE1 elements in WT and *Suv39h1KO* cells at steady state. Summary data are depicted as histograms at the top of the read alignment. (C) Pie chart showing percent enrichment of LINE1 subfamilies marked by *Suv39h1-*dependent H3K9me3 peaks displayed in B. Percent enrichment is indicated by emboldened numbers as in panel A. (D) Bar chart showing the percentage of total *Suv39h1*-dependent H3K9me3 peaks lost during LPS challenge. (E) Aligned *Suv39h1*-dependent LINE1 H3K9me3 peaks in untreated, two, and four hours LPS treated WT BMDCs. (F) Pie chart showing percent enrichment of LINE1 subfamilies in the LPS-sensitive H3K9me3 peaks on LINE1 in panel E. Percent enrichment is indicated by emboldened numbers as in panel A. (A-F) WT n=3, KO n=3 mice per experimental condition. In panels B and E, the x-axis coordinates refer to kilobases. Statistical significance in panel A was determined by a two-proportions z-test for comparison of proportions in two independent samples. Standard statistical analysis were used for peak calling of ChIPseq data (see materials and methods for detailed protocol.

### Innate activation regulates TE heterochromatin dynamics

Given that *Suv39h1* was uniquely depleted at both the mRNA and protein levels we sought to establish the overlap between enhanced TE expression and *Suv39h1*-dependent, LPS-sensitive, H3K9me3 peaks. H3K9me3 chromatin immunoprecipitation followed by high throughput sequencing (ChIP-seq) of WT and *Suv39h1*KO BMDCs allowed us to establish *Suv39h1*-dependent H3K9me3 patterns. Our analysis identified H3K9me3-bound regions, with approximately 80% overlap of H3K9me3 peaks among biological replicates of the same time point and background (Figure S2A). Consistent with the demonstrated role for *Suv39h1* in establishing H3K9me3 marks on TEs (*27*), we observed 126,000 H3K9me3 peaks at TE loci in *Suv39h1*WT reduced to 92,000 in *Suv39h1*KO BMDCs (Figure S2B). We reasoned that by filtering the WT BMDC H3K9me3 peaks for corresponding peaks in *Suv39h1*KO BMDC, then the remaining set of H3K9me3 peaks should represent those that are specifically regulated by *Suv39h1*. This analysis identified 75,000 peaks that we termed *Suv39h1*-dependent peaks (Figure S2B). Importantly, more peaks were identified as *Suv39h1*-dependent than were apparently lost in *Suv39h1*KO cells (Figure S2B, see methods) because our analysis included some peaks which were found in proximal but distinct positions in *Suv39h1*KO cells. We attribute these peaks to compensatory methylation by *Setdb1,* which normally targets a set of TEs that is broader but overlaps with *Suv39h1* targets (*19*). Altogether, our analyses suggested that *Suv39h1* maintained 58% of the H3K9me3 peaks in untreated WT cells (Figure 2A, S2B). Strikingly, when compared to *Suv39h1-*independent peaks, *Suv39h1* H3K9me3 peaks were significantly enriched in LINE1 (60%), whereas LTR (26%), and SINE (7%) were relatively depleted (Figure 2A). Of note, and consistent with the impact of *Suv39h1* knockout on TE expression (Figure S1D-F), *Suv39h1-*dependent peaks at LINE1 were primarily enriched on evolutionarily younger elements (L1Md_F, L1Md_T, L1Md_A) (Figure S2C). Peak inspection revealed robust and selective enrichment of H3K9me3 at the 5’ end and to a much lesser extent at the 3’end of intact LINE1 elements (Figure 2B). Deletion of *Suv39h1* abolished H3K9me3 at the flanking regions as well as within the bodies of LINE1 elements (Figure 2B, S2C). Although *Suv39h1-*dependent H3K9me3 was not especially enriched on SINE elements relative to *Suv39h1*-independent peaks (Figure 2A), a subset of peaks on SINEB1 were *Suv39h1-* dependent (Figure S2D). In contrast, the impact of *Suv39h1* deletion was minimal at other TEs such as SINEB2, ERV1, ERVL, and ERVK (Figure S2F), consistent with reduced enrichment of *Suv39h1-*dependent H3K9me3 peaks at these elements (Figure 2A). Again, we hypothesize that the reduced effect of *Suv39h1*KO on H3K9me3 at these elements may reflect compensatory methylation by *Setdb1* (*19, 35*).

Next, we treated WT BMDCs with LPS to compare *Suv39h1*-dependent and LPS-sensitive H3K9me3 peaks by ChIP-seq. Consistent with LPS-driven loss of *Suv39h1* expression, LPS stimulation of WT BMDCs readily extinguished H3K9me3 at specific loci, totaling 35% of *Suv39h1*-dependent H3K9me3 peaks (Figure 2D). Concordantly, these peaks were lost at LINE1 and SINEB1 (Figure 2E, S2F). Among LINE1 elements that lost *Suv39h1*-dependent H3K9me3 during LPS challenge (Figure 2E), the L1Md_F2 subfamily was disproportionately enriched (47%) compared to other LINE1 subfamilies (Figure 2F), indicating direct impact of *Suv39h1* loss on these elements during LPS treatment. In contrast, although LPS stimulated LTR element expression (Figure 1), it did not extinguish H3K9me3 peaks at ERV1 and ERVK elements as readily as at LINE1 and SINEB1 (Figure S2G), consistent with the reduced effect of *Suv39h1* knock-out on steady-state H3K9me3 at these loci (Figure S2E). We propose *Suv39h1*-independent regulation of these elements downstream of LPS. Our data support a model in which LPS-induced depletion of SUV39H1 (Figure 1G, H) facilitates heterochromatin loss to directly regulate de-repression of specific LINE1 and SINEB1 elements. This model is consistent with observations that disruption of epigenetic pathways that silence LINE1 causes their expression and activation of innate immunity (*13, 30, 31*). We hypothesize that specific TEs are regulated as part of normal innate immune physiology and might naturally contribute PAMPs to enhance, amplify, or broaden a physiological innate immune response.

### TE expression and inflammation

We next sought evidence for TE contributing endogenous PAMP that might be detected and therefore contribute to LPS-driven inflammatory responses. We first examined the effect of overexpressing *Suv39h1* on LPS treatment of RAW264.7 macrophages (Figure 3A). *Suv39h1*-overexpression (SUV39H1oe) significantly dampened the LPS-mediated induction of a luciferase construct controlled by the ISG54 promoter construct measured as a surrogate for IFN-I production and ISG induction (ISG54-luciferase). LPS induction of the endogenous ISG VIPERIN was also repressed by SUV39H1oe (Figure 3B). This effect was confirmed at the level of signal transduction with reduced levels of phosphor-TBK1 and -IRF3 but unaffected levels of total TBK1/IRF3 protein (Figure 3D), consistent with Suv39h1oe reducing PAMP levels and thus downstream PRR signaling. Importantly, SUV39H1oe did not impact ISG54-luciferase production after stimulation with a STING ligand, 2’3’cGAMP (Figure 3C), demonstrating that cells overexpressing *Suv39h1* were capable of producing WT levels of ISG activation downstream of DNA sensing activation.

**Figure 3.**
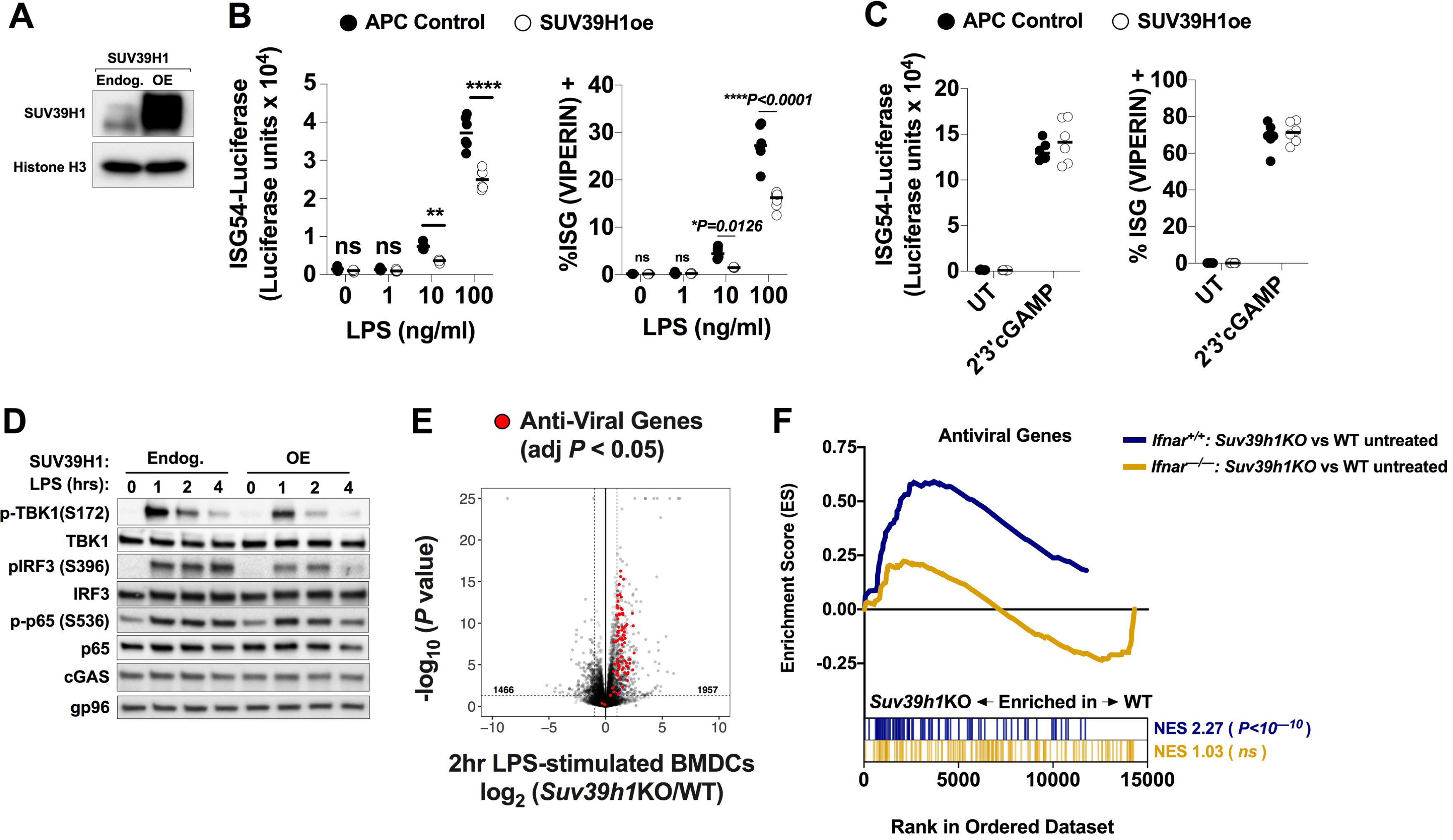
LPS-elicited physiological depletion of SUV39H1 promotes a type-I IFN response in myeloid cells independently of direct regulation of ISG loci, see also Figure S3. (A) Western blot of SUV39H1 in RAW264.7 macrophages transduced with empty lentivector or with a SUV39H1 overexpression (OE) lentiviral vector. (B) Left, RAW264.7 macrophages transduced with lentivirus control (n=5, closed circles) or SUV39H1 overexpressing lentivirus (n=5, open circles) were stimulated for 8 hours with the indicated concentrations of LPS and the IFN-I response measured was measured by ISG54-driven luciferase assay. Right, flow cytometric analysis of intracellular expression of the ISG, VIPERIN, by APC-control and SUV39H1 overexpressing RAW264.7 marophages stimulated as in left panel. (C) Left, RAW264.7 macrophages transduced with lentivirus as in B and stimulated with the STING ligand 2’3’cGAMP for 8hrs and the IFN-I response measured by ISG54-driven luciferase assay as in B. Right, flow cytometric analysis of ISG, VIPERIN, expression in untreated and 2’3’cGAMP transfected cells after 8hrs. (D) Western blot analysis of innate signal transduction pathways elicited in control and SUV39H1 overexpressing RAW264.7 macrophages by LPS stimulation for the indicated times. (E) Volcano plot depicting differential gene expression in WT versus *Suv39h1*KO BMDCs at two hours of LPS stimulation. anti-viral genes with an adjusted *P* value <0.05 and FC ≥ 2 are indicated in red. (F) Gene set enrichment analysis (GSEA) comparing expression of anti-viral genes from mRNAseq data between the following groups: untreated *Suv39h1*KO x *Ifnar^+/+^* (n=4) versus *Suv39h1*WT x *Ifnar^+/+^* (n=3) BMDCs (Blue); untreated *Ifnar^—/—^Suv39h1*KO (n=3) versus *Ifnar^—/—^ x Suv39h1*WT BMDCs (n=3) (gold). (D) Western blots of the indicated total proteins and phosphor-proteins during LPS treatment of BMDCs with the following genotypes from left to right: *Ifnar^+/+^*x*Suv39h1*WT*, Ifnar^+/+^*x*Suv39h1*KO*, Ifnar^—/—^*x*Suv39h1*WT and *Ifnar^—/—^*x *Suv39h1*KO. Western blots represent one of three experiments where similar results were obtained. In B and C, statistical significance was determined by two-way ANOVA and Bonferonni ad-hoc analysis: \**P*<0.05, ** *P* <0.001, \*\*\**P*<0.001, \*\*\*\**P*<0.0001. In (E), significance cutoff for differential gene expression analysis was defined as an adjusted *P*-value < 0.05 and a fold change ≥ 2. WT, n=3 mice per group; KO, n=4. In (F), statistical significance was GSEA; *Suv39h1*WT x *Ifnar^+/+^* (n=3)*, Suv39h1*KO x *Ifnar^+/+^* (n=4)*, Suv39h1*WT x *Ifnar^—/—^* (n=3), and *Suv39h1*KO x *Ifnar^—/—^* (n=3).

Conversely, complete ablation of *Suv39h1* had opposite effects. LPS-stimulated *Suv39h1*KO cells produced significantly higher levels of IFN-I (IFN) mRNA and protein than their WT counterparts (Figure S3A, B), exhibiting enhanced phosphor-TBK1, -p65, and IRF3 signal-transduction (Figure S3C). Concordantly, *Suv39h1*KO cells exhibited a more vigorous and sustained anti-viral response to LPS than WT cells, evidenced by the upregulation of ISGs (Figure 3E, S3D). We conclude that *Suv39h1* levels, up or down, determine the magnitude of the IFN-I/ISG response following LPS stimulation.

In parallel experiments, we found that *SUV39H1* depletion with shRNA in primary human monocyte-derived DCs also significantly enhanced LPS induced ISG SIGLEC-1 expression whereas it had less pronounced effects on expression of the NFkB-regulated activation marker CD86 (Figure S3E,F). Our data are therefore consistent with SUV39H1 regulation of LPS responses being conserved in human cells.

We hypothesized that enhanced steady-state expression of TEs in *Suv39h1*KO cells should also amplify baseline ISG expression. In fact, gene set enrichment analysis (GSEA) of RNAseq data revealed that *Suv39h1*KO BMDCs exhibited a significantly augmented type-I IFN/anti-viral gene signature at baseline compared to WT cells (*P < 10^-10^*) (Figure 3F, blue). Further, ablation of the type-I IFN receptor (*Ifnar*) in WT and *Suv39h1*KO BMDCs, abolished baseline differences in ISG expression (*P > 0.05*) (Figure 3F, gold), confirming that these ISG steady-state differences resulted from enhanced type-I IFN signaling. In a second cell type, in RAW264.7 macrophages, depletion of *Suv39h1* with shRNA (Figure S3G, H) also enhanced the baseline ISG expression as measured by ISG54-luciferase production (Figure S3I). These data suggest that *Suv39h1*KO is not directly impacting ISG expression because increased ISG expression levels in *Suv39h1*KO cells is IFN dependent.

To further probe whether ISG and cytokine loci are directly regulated by *Suv39h1* and H3K9me3, we analyzed H3K9me3 peaks in ISG loci and compared them directly with mRNAseq data from the same samples. H3K9me3 assessment by ChIPseq revealed that ISGs, and *Ifnb* itself, are not associated with Suv39h1-sensitive H3K9me3 marks (Figure S3J). Furthermore, both WT and KO cells broadly maintained H3K9me3 peaks in the same set of ISGs, and in fact, *Suv39h1*KO cells maintained slightly more peaks (Figure S3J). In addition, the presence or absence of H3K9me3 peaks did not correlate with ISG mRNA expression at baseline (Figure S3J). Therefore, direct regulation of ISGs by *Suv39h1* and H3K9me3 does not explain the enhanced ISG response in *Suv39h1*KO cells. Rather, our data suggest that *Suv39h1* loss, either downstream of LPS signaling, or due to *Suv39h1*KO, drives TE expression which activates innate immune sensing and IFN production.

### cGAS is required for *Suv39h1*-mediated effects on IFN-I

We hypothesized that LPS-induced TE expression is detected by innate sensors thereby contributing endogenous PAMP to LPS responses. Nucleic acid sensors are thought to detect TE-derived PAMPs when TE are expressed in various experimental systems: knock out of epigenetic regulators (*13*), models of senescence/aging (*30, 31*), cancer (*9, 36*). Furthermore, in the autoinflammatory condition, Aicardes Goutierres syndrome (AGS), individuals defective for the nuclease TREX1, which ordinarily degrades cytoplasmic DNA, experience cGAS-dependent (*37*) IFN-I-driven, inflammation resulting from either chromatin leakage into the cytosol or from TE-derived DNA (*38, 39*) with the contribution of each still debated. It is unclear whether TEs contribute PAMP to natural, physiological innate immune sensing.

We therefore explored the contribution of *Suv39h1*-regulated TE expression to LPS responses. We first assessed activation of the DNA-sensing cGAS pathway by measuring STING phosphorylation (Konno et al., 2013) in LPS activated WT BMDCs by immunoblot. In this experiment, we increased the number of cells loaded into gel wells (8.5x10^5^) and transfected a low dose of 2’3’cGAMP to allow comparison of phosphor-STING as a control. Strikingly, LPS induced STING phosphorylation and this was lost in *Cgas*^—/—^ cells, consistent with endogenous PAMP detection by cGAS (Figure 4A). LPS-activation of IRF3 detected by WB was only modestly reduced by cGAS knock out, consistent with cGAS-independent LPS-induced canonical TRIF and MYD88 activation of IRF3 (Figure 4A). IRF3 activation by phosphorylation is a complex process, and one possibility is that cGAS impacts IRF3 phosphorylation at sites that are important for activation but that are not detected here (Robitaille et al., 2016). As expected, transfection of BMDCs with 2’3’cGAMP (250-500ng/ml) induced STING and IRF3 phosphorylation independently of cGAS (Figure 4A). Concordantly, LPS stimulation of *Cgas^—/—^* BMDCs induced significantly less IFNB (type I IFN) protein as compared to WT cells (Figure S4A). As a control, *Cgas* ablation blocked HT-DNA-induced IFNB whereas 2’3’cGAMP transfection induced similar IFNB levels in both WT and *Cgas^—/—^* cells as expected since 2’3’cGAMP activates STING directly (Figure S4B). Similar results were obtained using freshly isolated splenic DCs where LPS treatment of *Cgas^—/—^*cells also triggered reduced expression of the ISGs VIPERIN and CCL5 relative to WT cells (Figure S4B). Again, transfected 2’3’cGAMP induced normal ISG responses in both WT and *Cgas^—/—^* cells as expected. Importantly, surface expression of CD86, a NFkB-regulated gene, was not impacted by *Cgas*KO (Figure S4B).These data indicate a contribution of cGAS/STING DNA sensing in LPS-induced activation of the IFN-I/anti-viral pathway.

**Figure 4.**
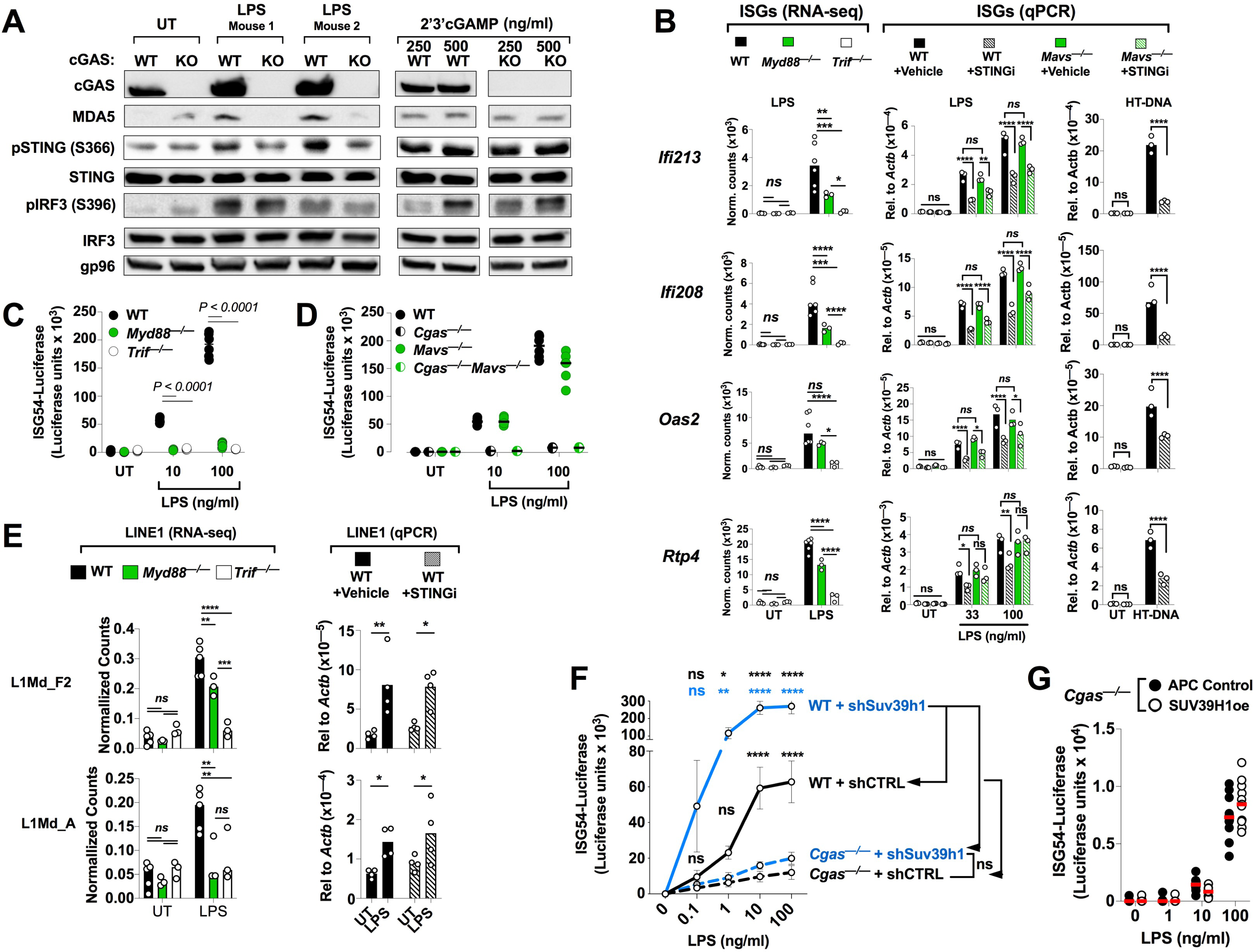
LPS elicits cGAS-dependent but not MAVS-dependent ISG expression, see also Figure S4. (A) Western blots of the indicated proteins and phosphor-proteins in WT and *Cgas^—/—^*BMDCs treated as follows: LPS treatment was for 4 hours; cGAMP (250-500ng/ml), 2 hours. (B) The indicated ISGs indicated in bold italics on the left were measured by mRNAseq of steady-state and LPS-stimulated (4hrs, 100ng/ml) WT, *Myd88^—/—^*, *Trif^—/—^* BMDCs or by RT-qPCR of WT and *Mavs^—/—^* BMDCs treated with STING inhibitor (STINGi, H-151) or vehicle control (DMSO) at steady state or stimulated with the indicated LPS dose (33 or 100ng/ml). As a positive control for inhibition of the cGAS/STING pathway, WT BMDCs were treated with STINGi or DMSO and then transfected with 1µg/ml HT-DNA for 5 hours. ISG expression was calculated relative b-Actin by the delta Ct method. (C) Scatter plot of ISG54-Luciferase assay of conditioned supernatants from WT, *Myd88^—/—^*, or *Trif^—/—^* RAW264.7 macrophges stimulated with the indicated concentration of LPS. (D) Scatter plot of ISG54-Luciferase assay of conditioned supernatants from WT, *Cgas^—/—^*, *Mavs^—/—^*, and *Cgas^—/—^ x Mavs^—/—^* RAW264.7 macrophages that were stimulated with the indicated concentration of LPS. (E) The indicated LINE1families indicated in bold on the left were measured by mRNAseq of steady-state and LPS-stimulated (4hrs, 100ng/ml) WT, *Myd88^—/—^*, *Trif^—/—^* BMDCs or by RT-qPCR of WT and *Mavs^—/—^* BMDCs treated with STING inhibitor (STINGi, H-151) or vehicle control (DMSO) at steady state or stimulated with the indicated LPS (100ng/ml) for 4hrs. As a positive control for inhibition of the cGAS/STING pathway. (F) ISG54-driven luciferase assay as in C and D. WT and *Cgas^—/—^* RAW264.7 macrophages in which *Suv39h1* was left intact (shCtrl) or depleted (*shSuv39h1*) with lentiviral-derived shRNA were treated with the indicated concentrations of LPS for 8. WT (*shCtrl*, n=8; *shSuv39h1*, n=8), *Cgas^—/—^*(*shCtrl*, n=8; *shSuv39h1*, n=8). (G) ISG54-driven luciferase assay as in C,D, and F. *Cgas^+/+^* or *Cgas^—/—^* RAW cells that were transduced with APC control or SUV39H1 overexpression lentivectors and stimulated with the indicated concentrations of LPS for 8hrs (n=10 per group). B-G, Statistical significance was determined by two-way ANOVA with Bonferonni’s ad hoc analysis for individual comparisons:\**P*<0.05, ** *P* <0.001, \*\*\**P*<0.001, \*\*\*\**P*<0.0001.

Interpretation of these experiments with *Cgas^—/—^* cells is somewhat complicated by previous observations of reduced baseline expression of ISGs in *Cgas^—/—^* cells (*40*). We reasoned that cGAS may constantly detect host/TE DNA, leading to low levels of IFN production, which prime innate immune responses in general, and that this effect is lost in *Cgas* knockouts. If this is the case, then ablation of tonic IFN-I signaling by *Ifnar* KO should eliminate the cGAS contribution to LPS responses. To investigate this possibility, we generated *Ifnar^—/—^* and *Cgas^—/—^Ifnar^—/—^* RAW264.7 macrophages with CRISPR-Cas9. Critically, LPS induction of IFNB was significantly compromised in *Cgas^—/—^Ifnar^—/—^* cells, as compared to *Ifnar^—/—^* cells (Figure S4G), demonstrating cGAS dependence in *Ifnar^—/—^* cells. We therefore conclude that the reduction of LPS-induced IFN-I production by *Cgas^—/—^* cells did not depend on the steady-state differences in baseline IFN signaling in *Cgas^—/—^*cells because we identified dependence on cGAS in *Ifnar^—/—^* cells. As controls, both WT and *Cgas^—/—^* RAW264.7 macrophages responded similarly to stimulation with IFNB by producing comparable levels of ISG54-luciferase (Figure S4H), demonstrating intact *Ifnar/Stat1* signaling that was effectively abrogated upon *Ifnar* deletion (Figure S4H). Further, LPS stimulation of *Ifnar^—/—^* RAW264.7 macrophages induced IFNB production (Figure S4G) but did not induce the ISG, VIPERIN (Figure S4I), as expected. We conclude that cGAS dependence of LPS-induced type-I IFN production is not due to reduced tonic IFN signaling in *Cgas^—/—^* cells.

To further probe the role of the cGAS pathway during LPS responses, we used a small molecule STING inhibitor (STINGi, H151) (*41*). Acute inhibition of STING permits assessment of the role of the cGAS/STING pathway in the absence of baseline effects. Strikingly, STING blockade significantly attenuated LPS-induced expression of specific ISGs (Figure 4B). As controls, *Myd88*KO in BMDCs dampened LPS-induced ISG responses while *Trif* ablation completely abolished them (Figure 4B), and H151 prevented activation of ISG expression after transfection with the cGAS ligand HT-DNA (Figure 4B), as expected. Similarly, LPS induction of the ISG, MDA5, measured by western blot was strongly reduced in *Cgas^—/—^* cells (Figure 4A). Unexpectedly, *Mavs* deletion had no effect on ISG induction by LPS (Figure 4B) whereas the responses to transfected PIC, or influenza A virus/Puerto Rico 8/1934/H1N1 (IAV/PR8) infection, were attenuated (Figure S4C,D). These results suggest that *Mavs*-dependent sensing of TE-derived RNA does not contribute to LPS responses.

We next sought to confirm these observations in RAW264.7 macrophages using CRISPR-Cas9 genetic ablation of *Mavs,* in WT or *Cgas^—/—^* cells(Figure S4E). LPS induction of ISG54-luciferase expression was dependent on *Myd88*, *Trif,* and *Cgas* because individual knock outs showed significant impairment of IFN-I activation, recapitulating our results in BMDCs (Figure 4C,D). Once again, inhibition of RNA sensing by deletion of *Mavs* in a WT or *Cgas^—/—^* background, did not affect the LPS-induced ISG response (Figure 4D). As a control, ablation of *Cgas* and *Mavs* effectively blocked induction of IFN-I following transfection with HT-DNA or PIC (polyI:C), respectively (Figure S4F). Collectively, these data verify that cGAS-dependent sensing contributes to LPS responses, but that MAVS-dependent RNA sensing does not.

Our data suggest a mechanism in which *Trif* and *Myd88-*dependent genes, LPS induces *Trif* and/or *Myd88*-dependent TE expression (Figure 1B-D, S1B, D-F) that can be detected by cGAS. Importantly, while deletion of *Trif* or *Myd88* significantly blocked TE induction, inhibition of STING with H151 left TE induction intact (Figure 4E). Thus TE induction is downstream of LPS, and cGAS/STING activation is downstream of TE expression. Together, these data are consistent with a model in which the defect in LPS-triggered ISG responses in *Cgas^—/—^* cells stems from an inability to sense TE-derived DNA. Based on this model, we hypothesized that the enhancement of IFN production by *Suv39h1* depletion required *Cgas.* Indeed, *Cgas* deletion in RAW264.7 macrophages blocked enhancement of LPS-induced IFN-I production by *Suv39h1* shRNA (Figure 4G). Notably, enhancement of the LPS-induced IFN-I response in *Suv39h1*-depleted cells was independent of *Ifnar*, confirming our observations in BMDCs, and dependent on *Cgas* (Figure S4I). Therefore, depletion of *Suv39h1* augments LPS-induced IFN in a *Cgas*-dependent manner and is independent of tonic IFN signaling.

Consistent with *Suv39h1-*knockdown experiments, SUV39H1oe did not suppress LPS-induced IFN-I when *Cgas* was ablated (Figure 4F, compare to Figure 3B, note the difference in scale). Importantly, SUV39H1oe did not impact cGAS protein levels (Figure 3D) ruling out a direct effect on cGAS expression. Furthermore, SUV39H1oe did not restrict induction of IFN when cGAS was bypassed by direct activation of STING with its ligand, 2’3’cGAMP (Figure 3C), consistent with a role for *Suv39h1* in suppressing expression of endogenous cGAS ligands. Thus, *Cgas* is required for *Suv39h1* regulation of the type-I IFN response to LPS. These data are consistent with a model in which cGAS detects *Myd88/Trif*-driven TE expression following the physiological depletion of *Suv39h1* by LPS detection.

### cGAS binds LINE1 elements in LPS stimulated cells

The impact of *Cgas*/*Sting* blockade on the LPS-dependent IFN-I response suggested that LPS induces expression of TE that are reverse transcribed into DNA and detected by cGAS. We therefore sought evidence for LPS-induced cytosolic DNA. We isolated cytosolic fractions by digitonin extraction (*42*) and retained nuclei for standardized quantification. Real time-qPCR analysis of RNAse-treated cytosolic fractions from BMDCs revealed that LPS significantly boosted cytoplasmic levels of LINE1 and ERVK DNA (Figure 5A). Furthermore, pretreatment with a reverse transcriptase inhibitor cocktail (emtricitabine, tenofovir, and nevirapine) effectively ablated the increased cytosolic DNA levels (Figure 5A) and restricted expression of the ISG VIPERIN (Figure S5A). Therefore, reverse-transcribed TE DNA is induced by LPS, and reverse transcriptase inhibition reduces LPS-induced ISG expression.

**Figure 5.**
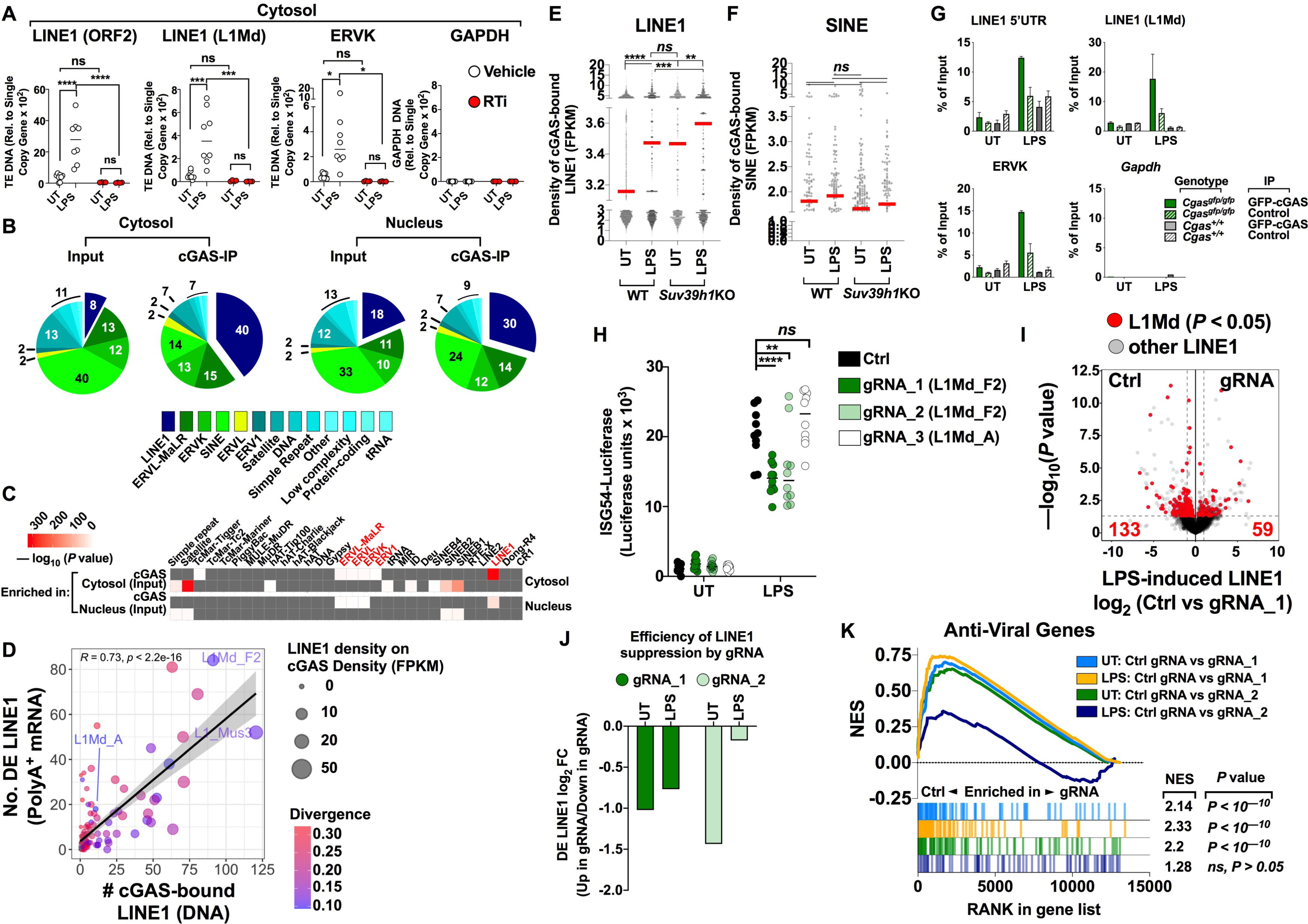
LINE1 is an endogenous cGAS PAMP, see also Figure S5. (A) qPCR of digitonin-extracted, RNAse-treated cytosolic extracts from untreated or LPS stimulated (100ng/ml, 4hrs) WT BMDCs that were pretreated with an RTI cocktail or DMSO vehicle control. (B) High-throughput sequencing of DNA extracted from the following fractions from LPS-treated BMDCs expressing *Cgas^gfp/gfp^*: cytosolic input fraction, cytosolic cGAS, nuclear input fraction, and nuclear cGAS. The pie charts display the enrichment in each fraction visualized as the distribution of the sum of the read densities (FPKM≥1) per unique element belonging to the indicated DNA species. N=2 for each sample. (C) Hypergeometric statistical comparison of read density proportions in panel B between input and the respective cGAS-IP within each cellular fraction (indicated in bold on right). Red color denotes —log_10_ (*P* value); grey, no statistical difference. (D) Correlation between the number of individual LINE1 elements within each subfamily expressed as mRNA and detected in the top 10% of fold changes in cytosolic cGAS-bound DNA. Dot size indicates the DNA density (FPKM) of cGAS-bound LINE1 families and the color scale indicates the average sequence divergence from consensus of the elements detected on cGAS. Less divergence indicates evolutionarily younger elements. (E) Scattered dot plots exhibiting FPKM of cGAS-bound unique LINE1 element DNA (FPKM ≥ 1 and FC ≥ 2 over input) in untreated and treated *Suv39h1*WT*xCgas^egfp/egfp^* and *Suv39h1*KO*xCgas^egfp/egfp^* BMDCs. Red bars represent median values; box ends represent quartiles. Statistical analysis was performed with one-way ANOVA followed by Bonferroni’s ad hoc analysis for individual comparison between samples. (F) Scattered dot plots as in E except FPKM values represent density of unique SINE element DNA bound to cGAS. Statistical analysis was performed as in E. (G) qPCR of LINE1, ERVK, and genomic *Gapdh* DNA bound to cytosolic cGAS immunoprecipitated with anti-GFP magnetic beads from BMDC cytosolic fractions as in B and E (filled green bars). Controls include control-bead IP from *Cgas^egfp/egfp^*(hashed green bars), GFP-bead (filled grey bars) and control-bead IP (hashed grey bars) from *Cgas^+/+^* BMDCs. (H) ISG54-Luciferase assay of resting and LPS-stimulated (8hrs) WT RAW264.7 cells expressing mutant d(dead)Cas9 KRAB fusion protein and control or LINE1-targeted gRNAs. (I) Representative volcano plot comparing differentially expressed (DE) LINE1 elements in gRNA_1 versus Ctrl cells stimulated with LPS. (I) Bar chart representing the log_2_ fold-change between DE LINE1 elements (Adjusted *P* value <0.05) in Ctrl (“Down in gRNA”) and gRNA-expressing cells (“Up in gRNA”) at steady state or after LPS treatment. (K) Above, GSEA of the effect of the specified gRNA versus control on anti-viral gene expression in resting and LPS-stimulated cells. Line color indicates the specific comparison. Below, gene rank plots. Each bar represents an anti-viral gene and its position along the x-axis indicates relative enrichment in the indicated group: left indicates enrichment in control cells; right, LINE1-targeted gRNA expressing cells. bar color corresponds to the same color used to specify the specific comparison above. *P* value to the right indicates statistical significance of enrichment of anti-viral gene expression in the control group versus LINE1-targeted gRNA-expressing cells. NES, normalized enrichment score. A, E, F, and H, statistical significance was determined by 2-way ANOVA, and Bonferonni’s ad hoc analysis was used for individual comparisons. A, E, F, H: \**P*<0.05, ** *P* <0.001, \*\*\**P*<0.001, \*\*\*\**P*<0.0001.

We next sought to understand the nature of the DNA activating cGAS during an LPS response using cross-linked immunoprecipitation of a tagged endogenous cGAS allele (eGFP-cGAS) from BMDCs (Gentili et al., 2019), followed by high through-put sequencing (HTS). We first biochemically fractionated BMDCs and confirmed efficient separation of the cytosol and nucleus by western blot and qPCR (Figure S5B,C). Western blots revealed cytosolic as well and nuclear cGAS protein (Figure S5B), as previously observed (*43–45*). Nuclear cGAS is not expected to sense chromatin (*45–50*), but it may sense de novo nuclear TE DNA, for instance. We therefore isolated endogenous cGAS-bound DNA from the cytosolic and nuclear compartments separately, after LPS stimulation, and subjected them to HTS. Because cGAS produces 2’3’cGAMP when bound to dsDNA, single-stranded DNA and RNA-DNA hybrids (*51–54*) we used an approach that would preserve all DNA species in the sequencing libraries (see materials and methods) to capture a global view of cGAS-bound nucleic acid. To avoid aberrant post-lysis binding between DNA and cGAS (*55*), we cross-linked the cells before lysis, ensuring exclusive detection of cytosolic DNA species bound to cGAS *in vivo*. Furthermore, after pull-down, the immunoprecipitates were stringently washed with detergent and high salt buffer, precluding a significant contribution for residual post-lysis binding events; cGAS-DNA interactions are expected to occur with weak affinity (Kd=10µM) (*56, 57*).

Inspection of all possible DNA species in the cGAS-bound fraction revealed striking differences relative to the cytosolic input (Figure 5B). Most notably, LINE1 DNA was profoundly enriched on cytosolic cGAS compared to input (5.1-fold, 40% versus 8%) (Figure 5B, C) whereas LTR families accumulated at much lower densities (Figure 5B, C). Conversely, cytosolic cGAS was depleted of satellite, SINE, and simple repeat DNA relative to input (Figure 5B,C). A similar trend was observed for the nuclear pool of cGAS although it did not approach the same magnitude of fold enrichment or statistical significance (Figure 5B,C). For example, LINE1 enriched weakly (1.68-fold, 30% versus 18%) on nuclear cGAS (Figure 5B), agreeing with a recent report (Gentili et al., 2019). Importantly, Figure 5C illustrates that significant enrichment of LINE1 on cytosolic cGAS cannot be explained by genomic frequency of LINE1 copies because the enrichment of LINE1 in the cytosolic input fraction (8%) is significantly below the expected genomic frequency (18%), and this is further supported by the statistical test used to compare cGAS-IP versus input (hypergeometric test) that takes into account genomic frequency.

To broaden our observations to a different cell type, we transduced RAW264.7 macrophages with lentivirus carrying eGFP-tagged cGAS and performed cross-linked IP from the cytosol followed by qPCR after LPS treatment (Figure S5D). Again, IP of cGAS from these cells revealed clear enrichment of LINE1 compared with non-TE DNA (Figure S5D) after LPS treatment. Although mitochondrial DNA has previously been shown to stimulate the cGAS/STING pathway (*58*), we did not detect appreciably enriched mitochondrial DNA on cGAS in either LPS treated BMDCs or RAW264.7 macrophages (Figure S5D,E), suggesting that mitochondria are not an important source of DNA PAMP downstream of LPS detection (mitochondria are not expected to be damaged by LPS exposure). We conclude that LINE1 is the primary endogenous ligand of cytosolic cGAS downstream of LPS stimulation. This is consistent with LPS-induced epigenetic derepression of LINE1 following depletion of the LPS-sensitive H3K9 methyltransferase, *Suv39h1*.

To better understand the contribution of the different cGAS fractions to the LPS response, we compared fold changes in binding densities of TE subfamilies for LPS-treated versus untreated cells in each fraction. The largest changes corresponded to strong increases in LINE1 and LTR binding densities on cGAS whereas we detected only modest changes associated with nuclear cGAS binding (Figure S5F), consistent with overall enrichment (Figure 5B, C). Enumeration of unique TE copies contributing to the largest fold-changes in the different fractions identified the LINE1 subfamilies L1Md_F2 and L1_Mus3 as the most highly enriched (Figure 5D), reflecting the specific regulation of these elements by *Suv39h1* (Figure 2F, S2E). Of note, these elements were among the youngest (Figure 5D, least diverged) and most highly expressed LINE1 elements in terms of mRNA (Figure S1C, Figure 5D), linking their availability for cGAS binding to transcriptional activation by LPS.

To confirm whether cGAS-dependent LPS-induced IFN-I requires direct cGAS-DNA binding and enzymatic activity, we reconstituted *Cgas^—/—^* RAW macrophages with WT cGAS (cGAS-FL), a cGAS DNA-binding mutant that lacks the DNA-binding N-terminal 160aa and harbors mutations in its zinc-thumb domain that prevent bound DNA from activating cGAS enzymatically (cGAS-DBM) (*59*), or a catalytically-dead cGAS mutant (cGAS-CD, E225A/D227A) that cannot produce 2’3’cGAMP (Kranzusch et al., 2013). Reconstitution levels of the WT and cGAS-mutants are shown in Figure S5G. Because cGAS-FL elevated IFN-I in untreated cells (Figure S5H), we subtracted baseline values from stimulation induced IFN-I following treatment with HT-DNA, 2’3’cGAMP, or LPS (Figure S5I). Critically, LPS-induced ISG54-luciferase expression was selectively restored by cGAS-FL, but not by either cGAS mutant (Figure S5I). Accordingly, HT-DNA transfection selectively activated cGAS signaling when cGAS-FL but not when cGAS-DBM or cGAS-CD were used (Figure S5I). In contrast, 2’3’cGAMP stimulated ISG54-luciferase in all settings, confirming that downstream signaling functioned normally (Figure S5I). These data reveal: 1) a specific requirement for cGAS during LPS stimulation and 2) that DNA-induced cGAS enzymatic activity is necessary for cGAS enhancement of LPS-triggered ISG responses.

Because *Suv39h1* regulates the magnitude and kinetics of LINE1 expression during LPS challenge, we predicted that *Suv39h1* deletion would enhance LINE1 DNA binding by cGAS. Indeed, LINE1 was detectable on cGAS at low levels in untreated WT cells, but LPS treatment significantly increased LINE1 density (Figure 5E). Furthermore, in *Suv39h1*KO BMDCs, steady-state LINE1 densities reached magnitudes similar to that found on cGAS in LPS-treated WT cells. As expected these were further enriched upon LPS stimulation (Figure 5E). In contrast, as a control, SINE DNA revealed disparate trends with no statistically supported differences (Figure 5F). As a control for our HTS data, cGAS-IP, followed by qPCR, detected LPS-induced enrichment of LINE1 and LTR (ERVK) DNA on cytosolic cGAS whereas genic DNA such as *Gapdh* was not detected (Figure 5G). Comparison of the densities of DNA elements bound to cGAS exhibited significant and selective enrichment of LINE1 in *Suv39h1*KO relative to WT cells (Figure S5J,K). Examination of LINE1 elements precipitating with cGAS revealed that *Suv39h1* KO boosted the same LINE1 subfamilies that bound cGAS with high frequency in WT cells after LPS stimulation (Figure 5F, S5L). Thus, *Suv39h1*-regulated LINE1 elements bind cGAS following LPS detection.

### LINE1 elements act as endogenous PAMPs

To provide further validation for our model, we sought evidence for the role of LINE1s as endogenous PAMPs that induce IFN-I by targeting them for epigenetic repression using a CRISPR dCas9-KRAB fusion construct. In this system, the mutant Cas9 (dCas9) binds DNA but is incapable of generating dsDNA breaks that would likely occur in excess when targeting numerous LINE1 elements and cause cell death (*60, 61*). The fused KRAB domain recruits epigenetic factors such as SETDB1, a H3K9me3 methyltransferase, driving heterochromatization and subsequent transcriptional silencing of the target region (*61*). As a positive control, we targeted the 5’ untranslated region of *Itgax*, the gene encoding the highly expressed integrin and myeloid marker, CD11b. Using two different guide RNAs (gRNAs), we succeeded in selectively repressing CD11b while maintaining MHC-II expression (Figure S5M).

We next designed gRNAs to target 20,000-110,000 LINE1 elements belonging to the L1Md family, and specifically enriched either at the L1Md_F2 sub-family (gRNA_1 and gRNA_2) that bound at high levels to cGAS after LPS treatment or, as a negative control, at L1Md_A elements (gRNA_3) which bound at low levels to cGAS (Figure S5N). Consistent with the notion that elements that bind cGAS at higher levels should contribute the largest impact to cGAS-dependent IFN-I production, gRNA_1 and gRNA_2 significantly impeded IFN-I production; gRNA_3, however, had no effect (Figure 5H). Importantly, differential analysis of mRNA-seq data (Ctrl versus gRNA-expressing RAW cells) confirmed specific repression of L1Md expression by gRNA_1 and gRNA_2 (Figure 5I, J). Of note, the efficiency of LINE1 suppression by gRNA_1 was similar in both LPS-treated and untreated cells (Figure 5J, dark green bars) whereas gRNA_2 suppressed LINE1 in untreated cells but lost efficiency upon LPS exposure (Figure 5J, light green bars). The effects on LINE1 suppression were mirrored in the IFN-I anti-viral response as revealed by GSEA of anti-viral gene expression (Figure 5K); whereas gRNA_1 effectively attenuated anti-viral gene expression in both untreated and LPS treated cells, gRNA_2 selectively suppressed steady state expression, suggesting that its repressive effect was limited to baseline TE expression and overcome by a strong LPS-induced transcriptional stimulus. These effects cannot be explained by direct suppression of anti-viral genes because neither gRNA_1 nor gRNA_2 target sequences were appreciably enriched in the proximity of ISG loci (Figure S5O). Therefore, LINE1 expression is required for optimal induction of IFN-I by LPS.

Critically, in *Cgas^—/—^* macrophages, the suppressive effect of gRNA_1 and gRNA_2 on IFN-I production was lost (Figure S5P), similar to *Suv39h1oe* in *Cgas^—/—^*cells (Figure 4F). We conclude that LPS-induced LINE1 expression acts through cGAS to augment the IFN-I/anti-viral response. This observation is consistent with L1Md elements acting as LPS-induced endogenous cGAS ligands that amplify the type-I IFN response.

### Transposable element expression restricts RNA virus infection

Our data support a model where innate immune responses proceed through two phases: 1) initial detection of pathogen-derived PAMP drives expression of cytokines, IFN-I, and TE through classical innate signaling pathways and 2) once TE expression reaches a critical threshold, cell autonomous detection of TE expression by cGAS/STING, and possibly other sensors, amplifies the anti-microbial response (Figure 6A). Whereas bacteria encode both RNA and DNA ligands, viral genomes typically encode one or the other, providing a useful tool to physiologically test our model. For example, various RNA viruses, including influenza, activate TLR4 (*21*), and although they do not contain DNA, they can, unexpectedly, activate and/or express inhibitors for the cGAS/STING pathway (*62, 63*). IAV infection has previously been shown to induce LINE1 expression (*64, 65*). We therefore tested the effect of manipulating the mechanism of innate immune amplification, that we have described, on IAV infection.

**Figure 6.**
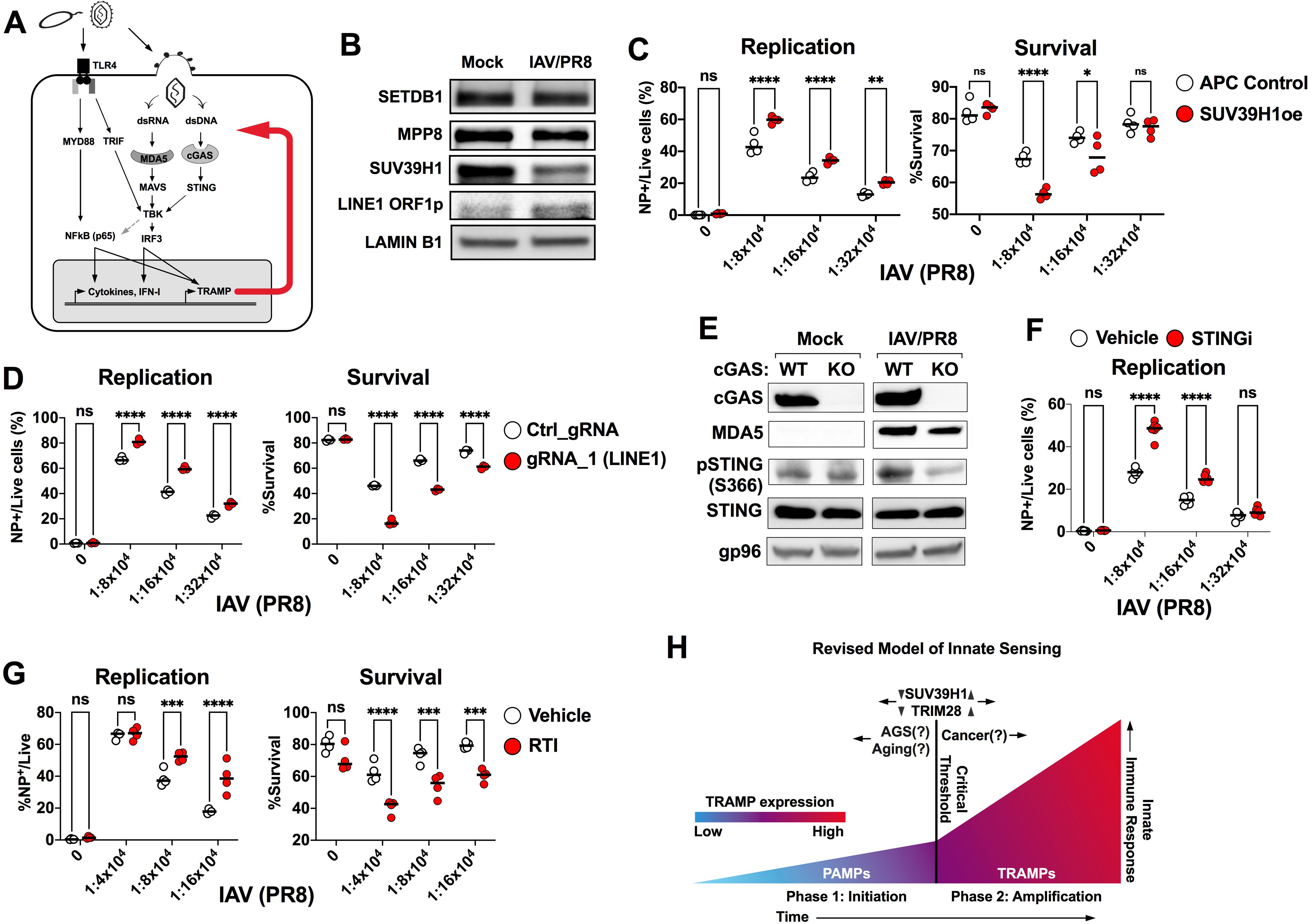
Induction of TE expression enhances host fitness during RNA virus infection. (A) Schematic illustrating the induction of endogenous PAMPs upon detection of microbial PAMPs. Microbial infection triggers IFN-I, cytokines and TEs. Subsequent detection of TE PAMP amplifies the innate immune response. (B) Western blots of the indicated proteins in in mock-treated or IAV/PR8-infected (7hrs) WT BMDCs. (C) Western blots of the indicated proteins and phosphor-proteins in mock-treated or IAV/PR8-infected (7hrs) WT and *Cgas^—/—^*BMDCs. (D) Quantification of IAV/PR8 NP expression in DCs 5 dpi in the presence of DMSO vehicle or STING inhibitor (STINGi, H-151). (E) DCs treated with DMSO vehicle or a RTI cocktail were infected with the indicated dilution of IAV/PR8 and NP expression(left) and percentage survival (right) were quantified by flow cytometry at 5dpi. (F) DC expressing lentivector control (APC) or overexpressing SUV39H1(oe) were infected with IAV/PR8 as in E and %survival and NP expression quantified 5dpi as in E. (G) DC expressing mutant d(dead)Cas9 KRAB fusion protein and control or LINE1-targeted gRNA_1 as described in figure 5 were infected with IAV/PR8 at the indicated dilution and NP expression(left) and percentage survival (right) quantified by flow cytometry. (H) Schematic displaying a revised mechanism for innate sensing of pathogen or that proceeds through two phases, in which TE expression is a central component. Initial detection of a pathogen occurs through classical sensing pathways that stimulate expression of defensive immune genes as well as TEs. Over time, TRAMP expression is amplified and may constitute the majority of detected PAMP in the latter phase. A critical threshold for TE detection may be surpassed once TRAMP concentration overcomes restriction factors such as the cytosolic DNase *Trex1,* for example. Grey arrow heads indicate increased or decreased expression; black arrows, impact on critical threshold; question marks indicate that the given condition is likely to impact the critical threshold but has not been directly tested. D-G, statistical significance was determined by 2-way ANOVA and Bonferonni’s ad hoc analysis for individual comparisons: \**P*<0.05, ** *P* <0.001, \*\*\**P*<0.001, \*\*\*\**P*<0.0001.

Srikingly, infection of WT BMDCs with IAV/PR8 selectively triggered depletion of SUV39H1 protein and induced expression of LINE1 ORF1p but did not appreciably affect SETDB1 or MPP8 levels at the time point queried (7 hrs post infection) (Figure6B). IAV infection can be assessed by cell survival and measurement of viral nucleoprotein (NP) (Figure S6). We therefore tested the physiological outcome of IAV/PR8-induced SUV39H1 depletion by measuring the impact of SUV39H1 overexpression on IAV/PR8 infection. Indeed, SUV39H1 overexpression rescued IAV infection, leading to reduced cell survival and higher levels of NP expression (Figure 6C), and consistent with the effect of SUV39H1 overexpression on cGAS-dependent LPS-induced IFN-I (Figure 3B, 4G). Next, we tested whether LINE1 suppression would rescue viral infection, hypothesizing that the depletion of SUV39H1 upon IAV/PR8 infection would promote innate detection of LINE1 PAMP, similar to LPS stimulation. Strikingly, suppression of LINE1 expression with dCas9-KRAB (Figure 6D) also rescued IAV/PR8 infection. We conclude that blockade of host LINE1 expression during viral infection reduced the murine innate immune response against IAV, leading to enhanced infection. Consistent with our model, we detected cGAS-dependent induction of phosphor-STING and the ISG MDA5 by IAV infection in DCs (Figure 6E). Furthermore, small molecule blockade of STING significantly increased IAV NP expression, consistent with STING signaling inhibiting RNA virus infection in these cells (Figure 6F). Concordantly, addition of a reverse transcriptase inhibitor cocktail, known to be active against TE including LINE1 (Banuelos-Sanchez et al., 2019), rescued IAV replication and enhanced virus-induced cell death (Figure 6G). Thus, consistent with our model, overexpression of SUV39H1, suppression of LINE1 expression, inhibition of STING, and inhibition of reverse transcription, all rescued IAV infection in DC through suppression of innate responses. These observations support our model that innate activation induces TE expression and reverse transcription-catalyzed TE DNA that is detected by cGAS to broaden and strengthen host defenses.

## DISCUSSION

In this study, we introduce a new paradigm for the mechanism of innate immune sensing in which endogenous PAMPs are produced within cells exposed to exogenous pathogen-derived PAMP. We propose that these endogenous PAMPs are detected by classical innate sensing pathways, amplifying and broadening responses to pathogens. Specifically, we discovered that LPS induces LINE1 expression, which is detected by cGAS to contribute to LPS-elicited IFN-I responses. We find that LPS activates LINE1 expression by driving SUV39H1 depletion that leads to loss of H3K9me3 at TE loci, combined with MYD88/TRIF-driven transcriptional activation. This discovery is important because it reveals an entirely new level of regulation of innate immune responses and ensuing inflammation that may explain inconsistencies in the field.

### Epigenetic regulation of TE in innate immune responses

The notion that TE should be permanently switched off derives from the observation that TE expression is typically associated with pathology, particularly autoinflammation and oncogenesis (*66–68*). We propose that our observation of regulated TE expression, and sensing by cGAS after LPS exposure, evidences an essential role for TE expression in normal innate immune responses. This is supported by our observation that disparate PAMPs also activate TE expression (Figure 1L).

Indeed, we hypothesize that diverse innate detection pathways converge on distinct epigenetic regulators to induce TE PAMPs that enhance innate immunity such as type-I IFN anti-viral responses.

Previous studies of TE regulation have suggested that expression of the youngest LINE1 is epigenetically regulated by an evolutionary Red Queen style arms race akin to that between host and pathogen (*20*). We hypothesize that such studies, characterizing the evolution of LINE1 regulation, demonstrate how host genomes have evolved to specifically regulate their expression when required. Strikingly, our data highlight the importance of LINE1 regulation in enhancing innate immunity and provide a direct example of the evolutionary benefits of retaining the capacity to regulate TE expression, rather than simply suppress it. Epigenetic marks such as H3K9me3 and DNA methylation switch gene expression on and off, and we propose that TE are simply another example of sequences that are epigenetically regulated in normal physiology, rather than comprising a distinct class of elements that must be permanently silenced (*19, 69*).

### DNA sensing of TE in innate immune responses

Our results highlight a central role for the DNA sensor cGAS in the LPS driven innate immune response. Interrogation of cGAS-bound DNA in LPS stimulated cells identified LINE1 DNA as a principal cytosolic cGAS ligand and our data demonstrating that reverse transcriptase inhibitors blocked LPS-induced cytosolic DNA, and that TE mRNA correlated with cGAS-bound DNA (Figure 5D), strongly implicate reverse transcription as a key step in PAMP generation. This was previously suggested in the AGS model, *Trex1^—/—^* mice (*70*). Indeed, RTI have been used successfully to treat AGS patients (*71*), reducing IFNa protein levels and ISG expression. In addition, in another study RTI treatment depleted LINE1 cDNA and senescence-associated inflammation (*30, 31*). Importantly, baseline expression of LINE1 RT, which promiscuously reverse transcribes host RNA, may generally prime innate immune sensing and IFN-I responses (*72–76*).

Our analyses indicated that the major changes in cGAS-bound DNA species during LPS stimulation occurred in the cytosol. The relative contribution of the nuclear and cytosolic cGAS pools to IFN-I production is not clear although it is clear that the nucleus potently restricts cGAS activation due to high affinity interactions with histones H2A and H2B (*45–50, 77*). Although nuclear cGAS is not expected to sense chromatin, it could conceivably sense nuclear de novo LINE1 DNA, per se. Indeed, LPS induced modest changes in binding between nuclear cGAS and TEs, but the majority of cGAS-TE interaction occurred in the cytoplasm. Moreover, condensation of cytosolic cGAS on DNA creates lipid droplets that can co-sediment with nuclei (Barnett et al., 2019; Du and Chen, 2018), suggesting that our observations of changes in the binding of nuclear cGAS with TEs are overestimated by cytoplasmic contamination.

Previous studies have demonstrated the activation of cGAS by mitochondrial DNA (mtDNA) (*58*). We ruled out a role for mtDNA, as it did not appreciably enrich on cGAS in either of the two cell systems interrogated here, BMDCs and RAW264.7 macrophages (Figure S5D, E). Thus, mtDNA binding to cGAS may be restricted by context such as certain types of viral infection (*78*) and pathogenic mutations in humans that permit leakage of mtDNA into the cytosol (*58*).

We were surprised to find in our experiments that RNA sensing of TE did not play a significant role in LPS responses because endogenous RNA PAMPs are expected to be produced more readily than DNA PAMPs which require reverse transcription from an RNA template.

Furthermore, RNA encoded by a ribosomal pseudogene has been implicated as an endogenous PAMP during infection, leading to a more potent anti-viral response against a variety of viruses including DNA viruses HSV-1 and EBV, and RNA virus IAV (*79*). It will be interesting to further investigate the impact of LPS on RNA sensing.

### Infection

We hypothesize that induction of endogenous PAMP after detection of exogenous PAMP is a powerful mechanism to broaden and amplify defensive responses. Indeed, a recent study showed that in a human lung epithelial cell line, IAV infection triggered sumoylation-dependent degradation of TRIM28, TE induction, and ultimately led to MAVS-dependent, STING-*independent,* sensing of, presumably, de-repressed ERV RNA (*64*). Furthermore, another publication suggested that a host commensal skin bacteria could induce ERV elements via activation of TLR2, and this triggered innate immune responses via cGAS/STING in keratinocytes that promoted adaptive T cell responses (*80*). Importantly, these responses were inhibited by reverse transcriptase inhibitors, indicating that reverse transcribed ERV cDNA was detected by cGAS and concurring with our data describing cGAS-mediated detection of reverse-transcribed LINE1 elements downstream of TLR4 signaling. These observations support our model that TE expression is an integral component of innate sensing that amplifies the primary innate response through subsequent cell-autonomous sensing events (Figure 6E).

Our model is entirely consistent with observations of viruses evolving antagonists for sensing pathways that they are not expected to activate. For instance, RNA viruses block DNA sensing pathways (*81*). Furthermore, RNA viruses such as SARS-CoV2 (*62*), measles, and Nipah virus (*82*) also trigger cGAS/STING sensing, and SARS-CoV2 at least has been demonstrated to induce LINE1 expression (*65, 83*). Concordantly, RNA viruses influenza (*84*), SARS-CoV (*85, 86*), SARS-CoV2 (*87*), Porcine epidemic diarrhea virus (*88*), Dengue virus (*89*), Yellow fever virus (*90*), Zika virus (*91*), and Hepatitis C virus (*92*) all inhibit the cGAS/STING DNA sensing pathway. Our model provides an explanation if detection of virus infection induces endogenous PAMP production. Consistent with this, we show that STING inhibition, reverse transcriptase inhibitors, LINE1 silencing, or overexpression of SUV39H1, all rescue IAV infection of mouse DC.

### Cancer

We propose that our model is applicable beyond infection. Natural TE expression correlates with better anti-tumor T cell responses (*93*) and the notion that enhanced TE expression drives an inflammatory response that is correlated with better clinical outcome, particularly after immunotherapy, is supported by studies that enhanced TE expression and the ensuing inflammatory responses either by small molecule inhibitors of DNA methylation or genetic ablation of epigenetic regulators(*9, 94–96*). Likewise, disruption of histone methylation by ablation of *SETDB1* led to TE dsRNA sensing and IFN-I responses in different cell lines including acute myeloid leukemia (*10*) and non-small-cell lung cancer(*11*). In addition, *Setdb1* deletion in a mouse tumor model led to enhanced T cell responses through presentation of TE-derived peptides rather than innate immune sensing (*97*). We speculate that a lack of innate recognition of TEs in this study may be linked to defective innate sensing pathways frequently encountered in cancer cells (Konno et al., 2018; Sutter et al., 2021). Based on our model, we hypothesize that high levels of TE expression and ensuing inflammation in tumors represents a natural innate immune defense against transformation. Concordantly, tumor cell intrinsic suppression of DNA sensing, for example via STING mutation, or suppression of TE induction by *Setdb1*, is common (*97–99*).

### Inflammation and Aging

The capacity of TEs to elicit innate immunity has also been linked with diseases other than cancer, particularly inflammatory disease. Patients with Aicardies Goutiere Syndrome (AGS) suffer an interferonopathy from early development which has been associated with TE expression, reverse transcription, and activation of cGAS (*37, 39, 71*). LINE1 DNA has also been revealed as a source of STING-dependent neuroinflammation in a *TREX1*-mutant model of AGS (*12*). To date, no associations have been made between the H3K9me3 pathway and AGS, although a recent paper linked diminished SUV39H1 expression with inflammation in the lungs of patients with chronic obstructive pulmonary disease (COPD) (*100*). Our data are consistent with these observations and may provide mechanistic explanation. TE expression has even been suggested as a driver of aging-associated inflammation (*30, 31*). Collectively, this body of literature provides numerous examples of TE-derived nucleic acids acting as endogenous PAMPs to activate innate sensors. However, the interpretation in each case has been that disease is enhanced or caused by the failure to keep TE expression switched off.

We propose, based on our observations, that the expression of TE to drive innate sensing is a natural physiological response to detection of pathogen and/or danger signals. Thus, disease stems from inappropriate regulation rather than simply failure to keep TE expression suppressed. For example, in cancer, we hypothesize that TE expression and activation of innate immunity is a response to transformation, evolved to activate adaptive immunity against transformed cells. We expect that the response of a cell, or cancer, to PAMP will vary depending on the active sensing pathways in each case. For example, many cell lines and cancers are defective for sensing (*98, 99*), and therefore, whether an RNA or DNA detection response dominates will depend on individual circumstances.

Boosting of an incipient innate immune response by autologous sensing of TE-derived PAMP is pivotal to our new model. To emphasize this and distinguish their role from that of conventional exogenous PAMP, we propose calling TE that are sensed during innate responses to infection <u>TRA</u>nsposable element <u>M</u>olecular <u>P</u>atterns (TRAMPs). In the two-phase mechanism (Figure 6H) that our data suggest, the initiating response, arising from direct recognition of exogenous PAMPs expressed by the infecting organism, activates expression of cytokines, ISGs, and, importantly, TRAMPs. In the amplifying phase, cell autonomous recognition of upregulated TRAMPs, by additional sensors including cGAS, amplifies the innate response. This model might explain how extremely low levels of exogenous viral PAMP such as in the case of natural HIV infection, for instance, can lead to a measurable inflammatory response. We imagine a threshold at which TRAMPs become dominant. This threshold can be shifted in disease states. For example, in AGS, the threshold is shifted to the left by, for example, defects that lead to increases in TRAMP levels (*TREX1*, *SAMHD1*). By contrast, in certain cancers, defects in nucleic acid sensing (cGAS, STING) shift the threshold to the right. Defects of epigenetic regulators could shift the threshold in either direction.

TE are not genetic junk left over from genome evolution. Rather, they are essential for the regulation of normal physiology. We anticipate that investigation of mutations and mechanisms regulating TRAMP expression will provide mechanistic insight into diverse drivers of inflammation in a wide range of conditions, including aging, cancer, and infection. This work could provide diagnostic tools and suggest novel therapeutic targets to transform our management of human disease.

---

**Figure S1.**
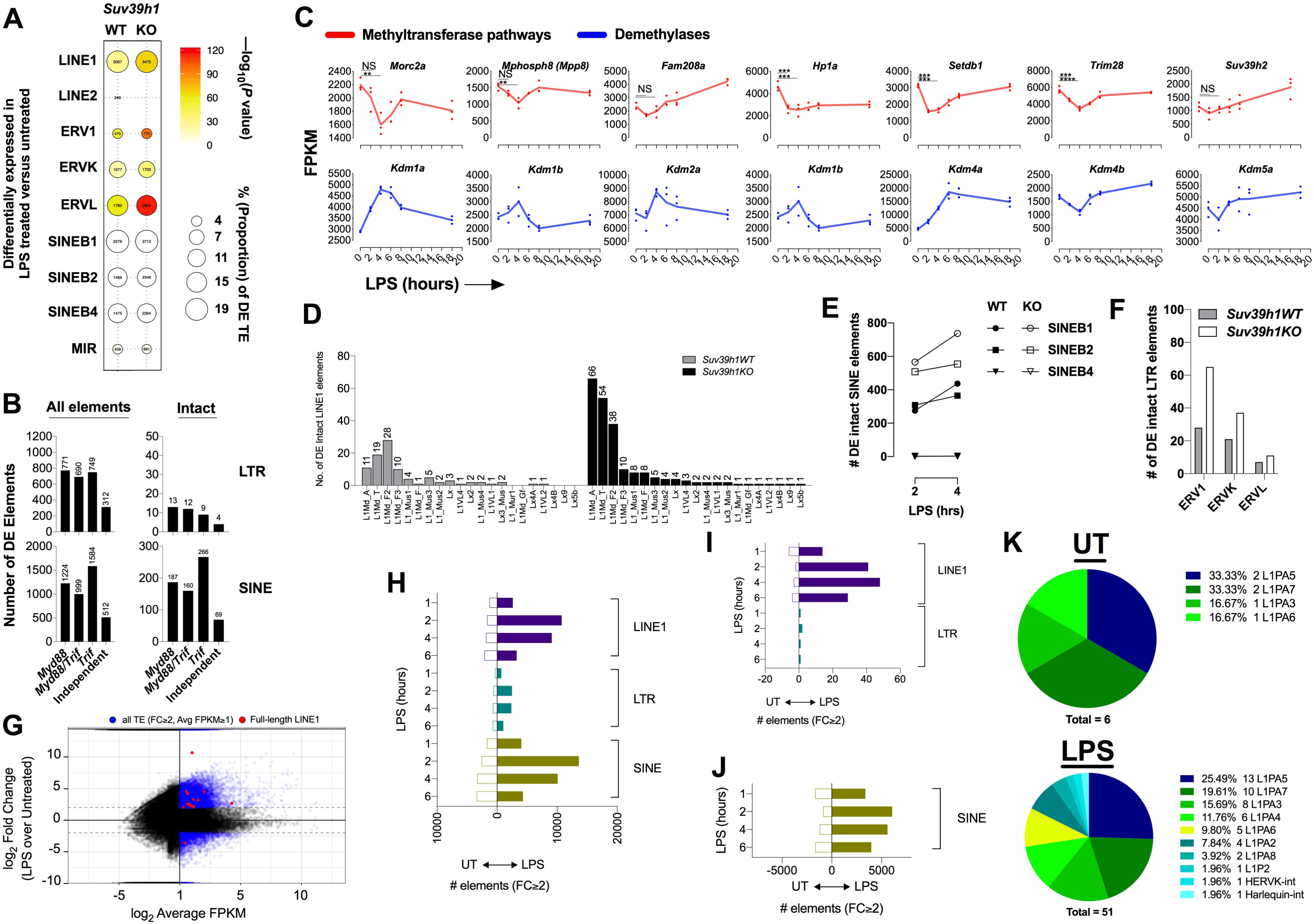
Ablation of *Suv39h1* significantly enhances LPS-induced intact TE transcription, reflecting loss of H3K9me3 during LPS challenge, related to Figure 2. (A) Bubble plot exhibiting percentage enrichment and statistical comparison of the proportion of differentially expressed TE in LPS treated versus untreated BMDCs in WT or *Suv39h1*KO cells. Statistical significance (P < 0.05) was determined by a two-proportions z-test. —log_10_ (*P* value) is indicated by color scale, bubble size indicates percentage of TE enrichment within each comparison. (B) mRNA sequencing data showing expression levels (FPKM) of H3K9me3 methyltransferases or pathway members (top, red lines) during LPS treatment: HUSH complex (*Fam208a, Mphosph8, Morc2a*) and other H3K9me3 remodelers/heterochromatin factors *Setdb1, Trim28, Suv39h2, and Hp1a (Cbx5)*. (Bottom, blue lines) mRNA sequencing data showing modulation of of histone lysine demethylases expression (FPKM): *Kdm1a/b, Kdm2a/b, Kdm4a/b*, and *Kdm5c*. (C-E) Summarized data of total numbers of intact LINE1 (C), SINE (D) and LTR (E) that were differentially induced in either *Suv39h1*WT or KO BMDCs when LPS treated samples were compared with steady state. (F) Mean average plot of human LINE, SINE, and LTR elements comparing 4hr LPS treated to untreated monocyte-derived DCs from human PBMCs. Blue dots, TEs with a log_2_FC 2, and average FPKM1; Red dots, full length LINE1 transcripts with a log_2_FC 2, and average FPKM1. (G) Enumeration of total TE numbers filtered by our criteria in panel e from each indicated class at the different time points of LPS stimulation compared to untreated cells. (H, I) Applying the same filters as in panel F, only the full length TE copies from each indicated class are displayed for LINE1 and LTR (H) and SINE (I). I, Pie charts exhibiting the relative proportions of expressed full length LINE1 copies enriched in untreated (UT, top) or LPS-treated (4hrs, bottom) cells.

**Figure S2.**
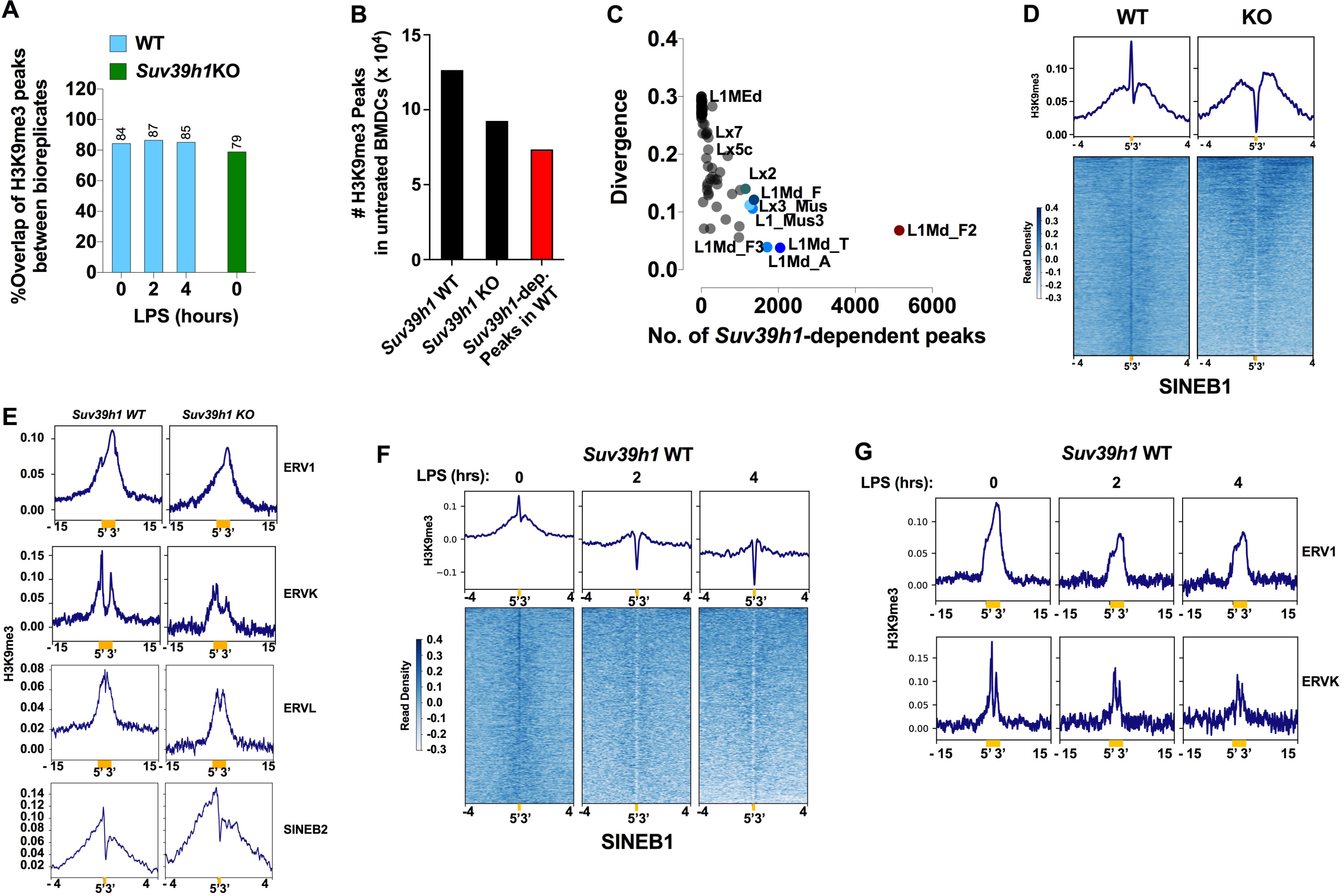
*Suv39h1*-dependent H3K9me3 peaks are primarily localized at transposable elements, related to Figure 1. (A) Percentage among three biological replicates of H3K9me3 peaks with significant overlap. (B) The number of H3K9me3 peaks in UT *Suv39h1WT* and *Suv39h1KO* BMDCs and the total number of *Suv39h1*-dependent peaks that were determined by subtraction of *Suv39h1*KO H3K9me3 peaks from WT H3K9me3 peaks. (C) Correlation between divergence of LINE1 versus the number of elements copies containing a *Suv39h1*-dependent H3K9me3 peak. (D) H3K9me3 peaks at SINEB1 elements in *Suv39h1*WT and *Suv39h1*KO BMDCs aligned at 5’ end with summary data at top as in Figure 2B, E. (E) Summary of H3K9me3 reads in the indicated elements that were aligned at their 5’ ends as in Figure 2B, E (top). (F) Aligned SINEB1 peaks as in D, in WT BMDCs stimulated with LPS for the indicated duration. (G) Summary data of aligned ERV1 (top) and ERVK (bottom) H3K9me3 peaks in WT cells stimulated with LPS for the indicated times. D-G, the x-axis coordinates refer to kilobases. Statistical peak calling was performed as detailed in the methods section.

**Figure S3.**
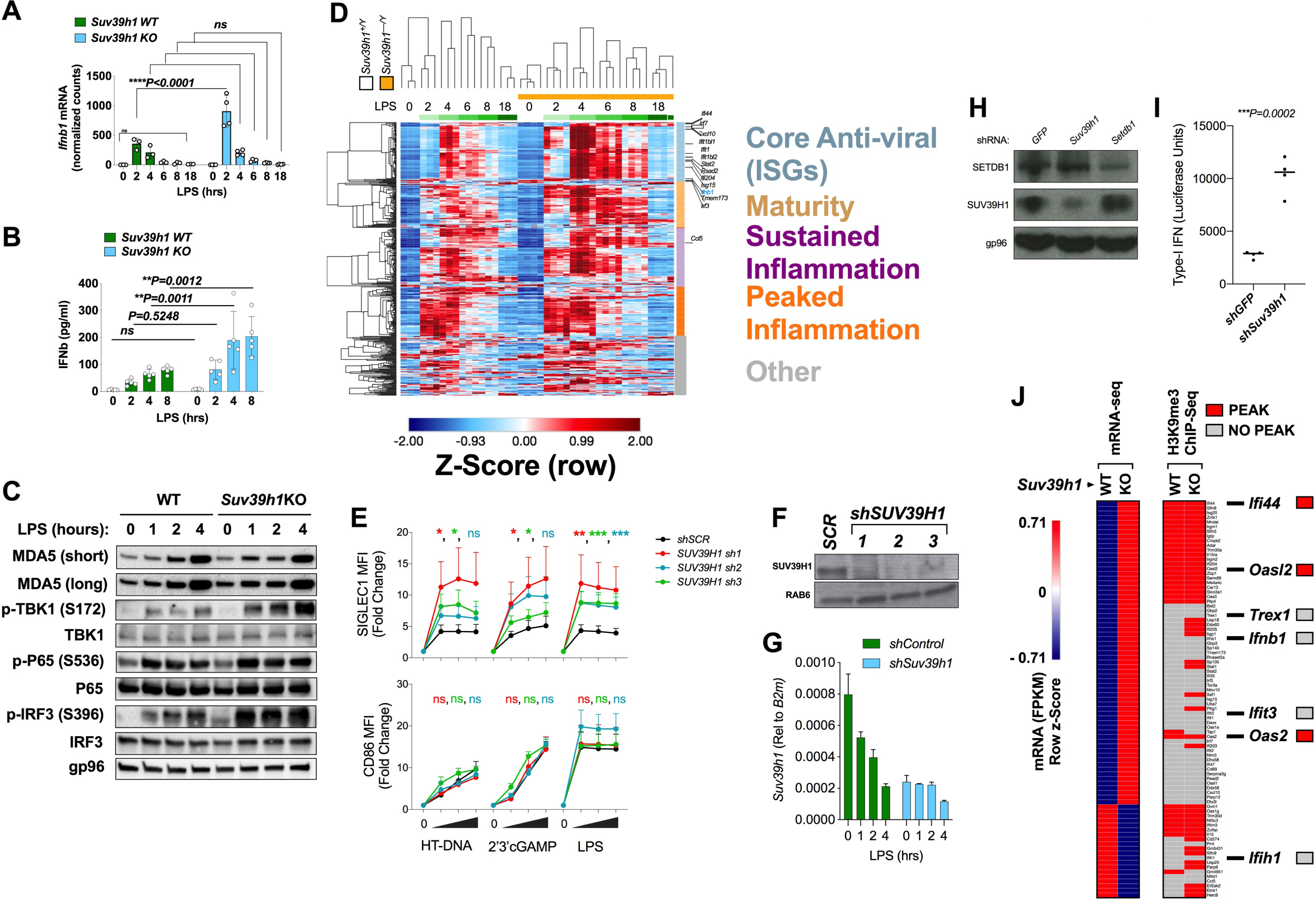
Loss of *Suv39h1* amplifies the LPS-induced anti-viral response independently of direct regulation of gene loci, related to Figure 3. (A) *Ifnb1* expression from mRNAseq data of *Suv39h1*WT and KO BMDCs that were treated with LPS for the indicated times. Data are displayed as FPKM. (B) ELISA quantification of Type-I IFN (IFNb) protein levels in conditioned supernatants from *Suv39h1WT* (n=4) or *Suv39h1KO* (n=5) BMDCs treated with LPS for the indicated times. (C) Western blots of the indicated proteins and phosphor-proteins in *Suv39h1*WT and KO BMDCs that were treated with LPS (100ng/ml) for the specified durations. (D) Heatmap displaying semi-supervised clustering (Pearson correlation) of anti-viral (ISG) and inflammatory gene expression in WT and *Suv39h1KO* BMDCs stimulated with LPS for the indicated times. Data are from the same mRNA-seq data set used for TE analysis in Fig. 1. (E) Flow cytometric data quantifying SIGLEC1 (ISG) and CD86 (NFkB-regulated) surface expression on mDCs transduced with nontargeting or one of three different SUV39H1-targeting lentiviral shRNA constructs (n=4 patient samples per condition). (F) Western blot of SUV39H1, demonstrating targeted depletion by three different lentiviral shRNAs in freshly cultivated human monocyte-derived DCs (mDCs). (G) qPCR data exhibiting the time course of *Suv39h1* mRNA expression during LPS treatment in RAW macrophages transduced with the designated lentiviral shRNAs (n=2). (H) Western blot of SUV39H1, SETDB1, and gp96 (loading control) to assess shRNA-mediated knockdown in untreated RAW264.7 macrophages transduced with the indicated lentiviral shRNA particle. (I) ISG54-Luciferase assay data comparing luciferase concentration in conditioned supernatants of untreated RAW264.7 in which *Suv39h1* was depleted by lentiviral encoded shRNA (*shSuv39h1*, n=4) or not (shGFP, n=4). (J) Left, heatmap showing z-scores of ISG mRNA expression (FPKM) in untreated *Suv39h1*WT (n=3) or *Suv39h1*KO (n=4) BMDCs, and right, the corresponding H3K9me3 peak status from H3K9me3 ChIPseq data (WT, n=3; KO, n=3): red, H3K9me3 peak positive, grey, no H3K9me3 peak detected. Several genes along with *Ifnb1* are designated in bold to the right as exemplars of ISGs with a grey or red box indicating H3K9me3 peak status. Statistical differences in panels A, B, and E were determined by two-way ANOVA, followed by Bonferroni’s ad hoc analysis for individual comparison between groups:\**P*<0.05, ** *P* <0.001, \*\*\**P*<0.001, \*\*\*\**P*<0.0001. In I, a two-tailed student’s t-test was performed to determine statistical significance (*P* < 0.05).

**Figure S4.**
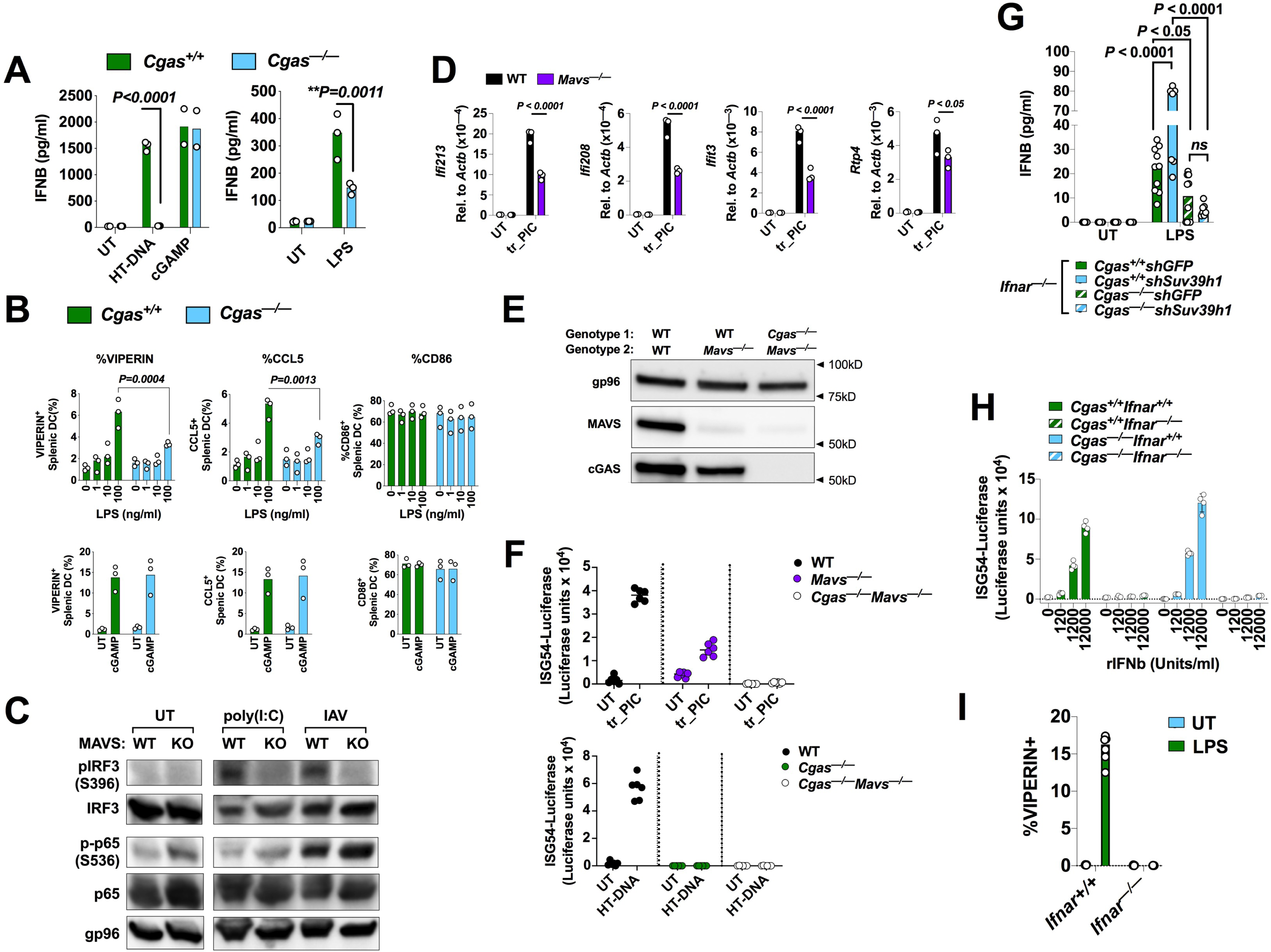
The cGAS pathway of DNA sensing, but not the RNA-sensing MAVS pathway is required for an optimal IFN-I/anti-viral response following stimulation with LPS, related to Figure 4. (A) ELISA quantification of IFNb protein concentration in conditioned supernatants from *Cgas^+/+^*(n=3) and *Cgas^—/—^* (n=3) BMDCs stimulated with LPS or with transfected HT-DNA, 2’3’cGAMP. (B) Compiled flow cytometric data showing the percentage of Viperin^+^ or CCL5^+^ CD11c^+^CD11b^+^MHC-II^+^ splenic *Cgas^+/+^* (n=3) and *Cgas^—/—^* (n=3) DCs stimulated *ex vivo* with 10-fold dilutions of LPS (top, 100ng/ml highest concentration) or bottom, with 2’3’cGAMP. (C) Western blot of the indicated total and phosphor-proteins following stimulation of WT and *Mavs^—/—^*BMDCs with transfected poly(I:C) (5hrs) or infection with IAV/PR8 (7hrs). (D) RT-qPCR data of the ISGs, 5hrs after transfection of poly(I:C) in WT and *Mavs^—/—^*BMDCs. ISG expression was calculated relative to b-Actin by the delta Ct method. (E) Western blot displaying protein expression of MAVS, cGAS, and gp96 after CRISPR-Cas9-targeted deletion of *Mavs* in WT and *Cgas^—/—^* RAW264.7 macrophages. (F) Functional validation of CRISPR-Cas9-targeted *Mavs* deletion in RAW264.7 macrophages displayed in E; top, transfection of poly(I:C), 2µg/ml; bottom, transfection of HT-DNA, 1µg/ml. (G) ELISA quantification of IFNB concentration in conditioned supernatants of untreated or LPS (100ng/ml) treated RAW264.7 macrophages that were deleted for *Ifnar1/2* by CRISPR-Cas9 and the transduced with control lentiviral particles (shGFP) or shRNA lentiviral particles targeting *Suv39h1 (shSuv39h1).* (H, I) Functional validation of CRISPR-Cas9 targeted deletion of *Ifnar* in RAW264.7 macrophges. (C) ISG54-driven luciferase assay measuring responsiveness of *Ifnar^+/+^Cgas^+/+^, Ifnar^+/+^Cgas^—/—^, Ifnar^—/—^ Cgas^+/+^,* and *Ifnar^—/—^Cgas^—/—^* cells to overnight stimulation with titrated recombinant IFNB protein.Error bars represent standard deviation and bar graphs represent the mean. Statistical significance was determined with two-way ANOVA and Bonferonni’s ad hoc analysis for individual comparisons. A, B, D, and G, statistical significance was determined with two-way ANOVA and Bonferonni’s ad hoc analysis for individual comparisons.

**Figure S5.**
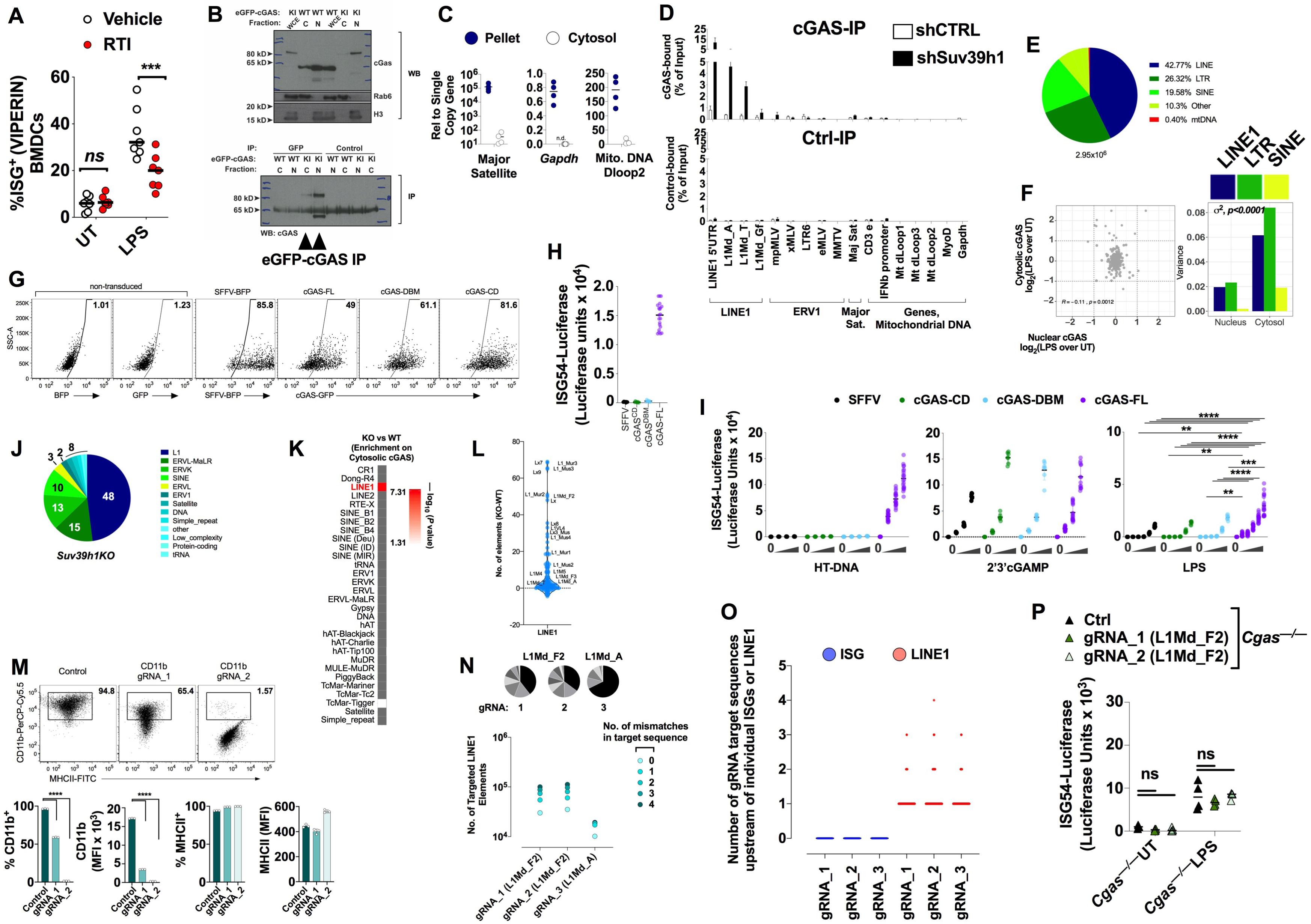
Immunoprecipitation of endogenous cGAS from nuclear and cytosolic extracts followed by NGS reveals significant changes in cytosolic cGAS-DNA binding activity, related to Figure 5. (A) Flow cytometric quantification of expression of the ISG VIPERIN in untreated or 8hr LPS-stimulated WT BMDCS that were pretreated with DMSO vehicle or an RTI cocktail. (B) Western blot revealing WT cGAS and the eGFP knock-in cGAS (KI) allele from whole cell lysate (WCE), cytosolic (C) and nuclear (N) fractions, as well as immunoprecipitation of cross-linked cytosolic and nuclear cGAS species. RAB6 and H3 serve as cytosolic and nuclear loading controls, respectively. Arrowheads indicate lanes displaying immunprecipitated eGFP-cGAS. (C) qPCR of the indicated amplicons in digitonin-extracted RNAse-treated cytosolic DNA and pellet fractions from untreated BMDCs. (D) qPCR of indicated DNA species that bound to an overexpressed eGFP-tagged cGAS contstruct in WT (shControl, n=4) and *Suv39h1* (n=4) knockdown RAW cells following LPS stimulation. (E) Pie chart depicting the relative proportions of raw reads in the indicated DNA species from NGS of cytosolic cGAS-IP. (F) Left, comparison of the fold changes in LPS versus untreated BMDCs of TE binding density on endogenous cGAS in cytosolic and nuclear fractions. Right, variance describing the fold changes in the binding density of the indicated TE class enriched on nuclear and cytosolic cGAS following LPS treatment (n=2 per condition). An F-test was used for statistical comparison of variances. (G) Flow cytometric plots displaying lentiviral-based expression levels of the indicated GFP-cGAS or SFFV-BFP control constructs used for experiments in Figure S5H, I. (H) ISG54-driven luciferase assay quantification of baseline IFN-I production by *Cgas^—/—^* RAW cells following lentiviral reconstitution with different cGAS constructs—full-length WT (cGAS-FL, n=12), catalytically-dead (cGAS CD, n=7), DNA binding mutant (cGAS-DBM, n=7), SFFV (empty vector control). (I) ISG54-driven luciferase assay quantification of IFN-I production by *Cgas^—/—^* RAW cells following lentiviral reconstitution with different cGAS constructs—full-length WT (cGAS-FL, n=12), catalytically-dead (cGAS-CD, n=7), DNA binding mutant (cGAS-DBM, n=7), SFFV (empty vector control)—and titrated stimulation with HT-DNA, 2’3’cGAMP, or LPS. Baseline values were subtracted from treated values to compare the response between groups reconstituted with distinct cGAS constructs. (J) Distribution of the cytosolic cGAS-IP read density sums (FPKM≥1) among all classes of DNA species in *Suv39h1KO* BMDCs. (K) Hypergeometric statistical comparison of the *Suv39h1*KO BMDC cytosolic cGAS-IP read density sums in panel E with WT cytosolic cGAS IP (Figure 5B). (L) Violin plot exhibiting the difference in the number of cytosolic cGAS-bound LINE1 elements by family between *Suv39h1*KO and *Suv39h1*WT BMDCs. (M) Inhibition of cell surface myeloid marker CD11b on RAW264.7 using the dCas9-KRAB transcriptional inhibitor system with two different gRNAs directed to the promoter region of the *Itgax* locus. Top, flow cytometric dot plots of CD11b (y-axis) and MHC Class II (MHC-II, x-axis); bottom, quantification of flow cytometric data for percentage of CD11b and MHC-II expressing cells and the MFI of each marker (n=3 per condition). (N) *In silico* prediction of the number of LINE1 gRNA targets (y-axis) according to the number of allowable mismatches between gRNA and target sequences (color intensity). Above, pie charts exhibiting enrichment of gRNA target sequences in LINE1 families. The most highly enriched family is denoted at the top of each pie chart and the corresponding gRNA number is listed below. (O) Scatter dot plot displaying the number of gRNA target sequences upstream of ISGs (blue circles) versus targeted LINE1 elements (red circles) . (P) ISG54-Luciferase assay of IFN-I and ISG induction in resting and LPS-stimulated (8hrs) *Cgas^—/—^* RAW264.7 cells expressing mutant dCas9 KRAB fusion protein and the indicated gRNA (Ctrl, gRNA_1, or gRNA_2; n=4 per group). Statistical significance was determined by 2-way ANOVA and Bonferonni’s ad hoc analysis for individual comparisons:\**P*<0.05, ** *P* <0.001, \*\*\**P*<0.001, \*\*\*\**P*<0.0001.

**Figure S6.**
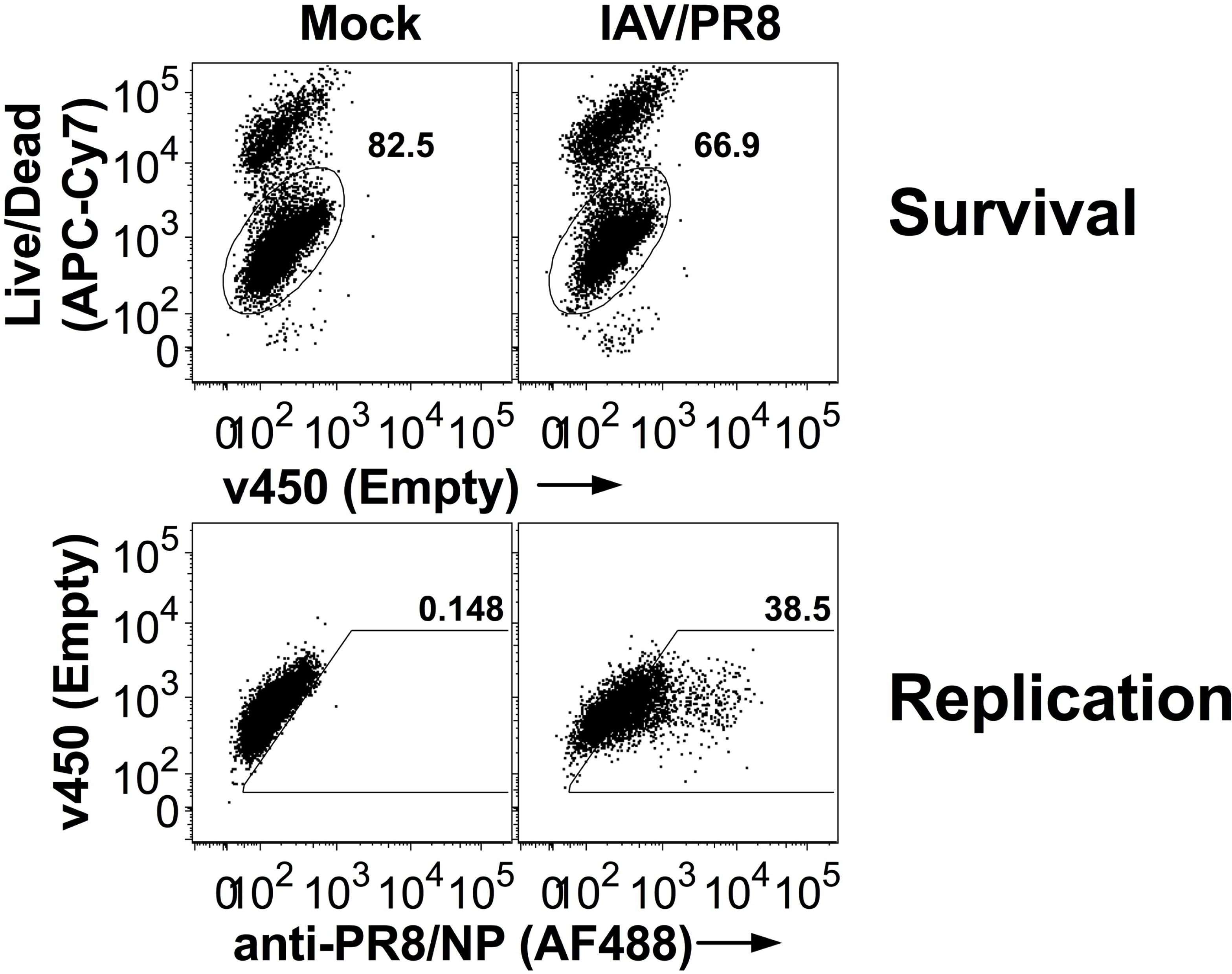
IAV/PR8 infection can be assessed by flow cytometric measurement of cell survival and viral nucleoprotein expression, related to figure 6. Top, flow cytometric dot plots of survival 5 days post infection (dpi) of DCs with IAV/PR8. Bottom, flow cytometric dot plots of IAV/PR8 NP protein expression 5 dpi with IAV/PR8 in the same sample as the corresponding plots above.

## ACKNOWLEDGMENTS

We thank J.M. Carpier, M. Burbage, M. Maurin, M. Gros, S. Heurtebise, L. Menger, A. Alloatti, and L. Joannas for assistance with various techniques and/or experiments. We thank R.E. Vance, J. Barau, L. Quadrana, and H. Rich for helpful discussions; M. Terman and G. Schwed, for helpful reading of the manuscript; S. A. Cros and S. Huertebise, for assistance with mice. We also thank the genomic and animal facilities. We thank T. Jenuwein for providing the *Suv39h1*-deficient mice. S. A. received funding from the Institute Curie; Institut National de la Santé et de la Recherche Médicale; Centre National de la Recherche Scientifique; ANR "ChromaTin" ANR-10-BLAN-1326-03, ANR “EPICURE” ANR-14-CE16-0009; S.A. received funding from la Ligue Contre le Cancer (Equipe labellisée Ligue, EL2014.LNCC/SA); Association de Recherche Contre le Cancer (ARC); grant ERC (2013-AdG N° 340046 DCBIOX); FP7_HEALTH-2010-259743 "MODHEP", ANR-11-LABX-0044_DEEP and ANR-10-IDEX-0001-02 PSL, ANR "CHAPINHIB" ANR-12-BSV5-0022-02, ANR "CELLECTCHIP" ANR-14-CE10-0013 and Aviesan-ITMO cancer project "Epigenomics of breast cancer". D.C.R was supported by funding from Institut National du Cancer (INCA) (grant 2017-1-PL BIO-05). High-throughput sequencing has been performed by the ICGex NGS platform of the Institut Curie supported by the grants ANR-10-EQPX-03 and ANR-10-INBS-09-08. N.M. and M.G. were supported by ANR (ANR-17-CE15-0025-01 and ANR-14-CE14-0004-02) and ERC (309848) grants to N.M. GK is supported by the Ligue contre le Cancer (équipe labellisée); Agence National de la Recherche (ANR) – Projets blancs; AMMICa US23/CNRS UMS3655; Association pour la recherche sur le cancer (ARC); Association “Ruban Rose”; Cancéropôle Ile-de-France; Fondation pour la Recherche Médicale (FRM); a donation by Elior; Equipex Onco-Pheno-Screen; European Joint Programme on Rare Diseases (EJPRD); Gustave Roussy Odyssea, the European Union Horizon 2020 Projects Oncobiome and Crimson; Fondation Carrefour; Institut National du Cancer (INCa); Inserm (HTE); Institut Universitaire de France; LabEx Immuno-Oncology (ANR-18-IDEX-0001); the Leducq Foundation; the RHU Torino Lumière; Seerave Foundation; SIRIC Stratified Oncology Cell DNA Repair and Tumor Immune Elimination (SOCRATE); and SIRIC Cancer Research and Personalized Medicine (CARPEM). This study contributes to the IdEx Université de Paris ANR-18-IDEX-0001.

## AUTHOR CONTRIBUTIONS

D.C.R. conceptualized, established, and performed histone ChIP-seq, cGAS-IP-seq protocols, cGAS-IP-seq library preparation, cell stimulations, cytosolic DNA extraction, CRISPR-dCas9-targeted suppression of LINE1 elements, lentiviral knockdown of *Suv39h1*, cGAS reconstitution, and SUV39H1 overexpression, IFNB and cytokine measurements, RNA-seq, and FACS analysis. M.G. made eGFP-cGAS lentiviral constructs and performed shRNA-targeted knockdown of SUV39H1 and stimulations in primary human MDDCs. N.B. and T.H. assisted D.C.R. with various experiments. P.B. performed analysis of RNAseq data from MDDCs, BMDCs and CRISPR-Cas9-KRAB suppression of LINE1. C.G. and M.Y. performed statistical analyses and mapping of H3K9me3-ChIP, cGAS-IP sequencing, and mRNA-seq of Suv39h1WT and KO BMDCs. N.M. supervised M.G. and contributed mice with an eGFP-tagged endogenous cGAS allele. D.C.R, P.B., M.Y., and C.G. analyzed data. D.C.R., G.J.T., and R.M. wrote the manuscript.

## DECLARATION OF INTERESTS

The authors declare no competing interests.

## MATERIALS AND METHODS

### Mice

*Suv39h1*KO(Peters et al., 2001) mice were from Thomas Jenuwein. *Mb21d1* (*Cgas*)^—/—^ (*B6(C)-Mb21d1^tm1d(EUCOMM)Hmgu^/J*) and *Mavs^—/—^* (B6;129-*Mavs^tm 1 Zjc^* /J) mice were from Jackson laboratory. cGAS-eGFP-Knock-in mice, a gift from Nicolas Manel (*Mb21d1^(tm1Ciphe)^;* CIPHE) were generated by targeted in-frame insertion of an enhanced GFP sequence after the ATG start codon of the endogenous *Mb21d1* locus. *Trif^—/—^*(C57BL/6J*-Ticam1^Lps2/J^*) and *Myd88^—/—^* (B6.129P2(SJL)-*Myd88^tm1.1Defr^/J*) were from Catherine Werts (Pasteur Institute) and *Ifnar^—/—^* (B6.129S2-*Ifnar1^tm1Agt^/Mmjax*) mice were originally from JAX. C57BL/6N mice were originally from Charles Rivers Laboratories. All mice were housed in SPF animal facilities, and live animal experiments were performed in accordance with the European Veterinary Department (Project Authorization N:02465.02).

### Cell Culture

Bone marrow-derived dendritic cells were cultivated in 20ng/ml GMCSF (Miltenyi) in IMDM (VWRI3390) supplemented with 10% fetal bovine serum (Eurobio), Penicillin/Streptomycin, 50µM -mercaptoethanol, minimal non-essential amino acids, and 2mM Glutamax (all from Life Technologies) and referred to as I-10 medium. Briefly, fresh bone marrow was collected from two of each—ilium, femur, and tibia—by centrifugation. Five million bone marrow cells were seeded on untreated 10cm plates (VWR) in 10mls of I-10 medium. On day 3, an additional 10mls of I-10 medium was added, followed by collection and replenishment of 10 mls on day six and on day eight if needed for use at day ten. BMDC clusters were harvested on day 8-10 in 5 mls of comlete I-10 media at room temperature and then stimulated at 1-2x10^6^ cells per well of an untreated 6-well plate (Sigma M9062-100EA) in 2 mls of I-10 medium without GMCSF.

### Cell stimulations

Cell stimulations were performed for the indicated times with mammalian cell culture-derived mouse IFNB (PBL, 12405), herring testes (HT)-DNA (1-4µg/ml as indicated; Sigma D6898-250MG), LPS (100ng/ml, Invivogen), high molecular weight poly (I:C) (1-2µg/ml, Invivogen), or 2’-3’cGAMP (4µg/ml, Invivogen) for the specified times. For cytosolic delivery of HT-DNA, poly(I:C), or 2’3’cGAMP, 12µg of PAMP and 12µl of Lipofectamine^TM^ (Life Technologies, 11668019) were separately diluted in OptiMEM (ThermoFisher 3195062), incubated for 5 minutes at room temperature, then combined, mixed by pipetting, and rested for an additional 15 minutes at RT. Liposome-containing PAMPs were then diluted in complete medium to make a concentrated working stock for cell stimulation. STING inhibitor (H-151, MedChem Express) stock solution was stored in DMSO at -80°C. Cells were pretreated with H-151 or DMSO vehicle for 2hrs before subsequent stimulation with PAMP or infection with IAV/PR8.

RAW264.7-ISG-Lucia cell lines (all from Invivogen) deleted for *Tmem173*(rawl-kostg*), Mb21d1*(rawl-kocgas*), Mda5* (rawl-komda5)*, and Rig-I*(rawl-korigi) carried a Lucia luciferase reporter construct under control of an IFN-inducible ISG54 promoter (Invivogen) and were cultured in DMEM containing 10% Serum (Fischer Scientific), 1% Penicilin/Streptomycin (Fischer Scientific) and Zeocin (1µg/ml; Invivogen). Cell stimulations were performed for the indicated times with HT-DNA, LPS, poly(I:C), or 2’-3’cGAMP, at the indicated concentrations in 6-well or 96-well tissue culture treated plates (Costar) following the same protocol as for BMDCs.

### Dendritic cell infection wit IAV

IAV/PR8/34 H1N1 was purchased from Charles River Laboratories and was stored at -80°C. Both BMDCs (Figure 6C) or the previously described JAWS II (ATCC)(*101*) mouse bone marrow-derived dendritic cell line (Figure 6B, D-G) was used for infection experiments. Briefly, BMDCs or JAWS II cells were plated in serum-free IMDM media supplemented with non-essential amino acids and infected with IAV/PR8 for 1hour at 37°C, before addition of complete media with a final concentration of 10%FBS. Infections were allowed to proceed for the times specified in figure legends. For RTI, treatment, emtricitabine and disoproxil fumarate were administered two hours prior to innoculation and maintained at 10µM for the duration infection (120hrs). DMSO vehicle control was administered alongside RTI treatment as a control.

### Generation of *Ifnar^—/—^* and *Mavs^—/—^* RAW264.7 macrophage cell lines with CRISPR

WT RAW264.7 macrophages were deleted using recombinant Cas9 protein (IDT, Alt-R S.p. HiFi Cas9 Nuclease V3) complexed with Tracr RNA and gRNAs targeting either *Ifnar1* (IDT pre-design gRNA Mm.Cas9.IFNAR1.1AA, Mm.Cas9.IFNAR1.1AB), *Ifnar2* (IDT pre-design gRNA, Mm.Cas9.IFNAR2.1AA, Mm.Cas9.IFNAR2.1AB), or *Mavs* (IDT pre-design gRNA, Mm.Cas9.Mavs.1AA, Mm.Cas9.Mavs.1AL). Briefly, gRNAs and Tracr RNA duplexes were generated by melting at 95°C for 5 minutes followed by downramping to 20°C at 5°C/minute. Duplexed RNA was then combined with recombinant Cas9 in duplex buffer for 15 minutes at room temperature to form Cas9 ribonucleoproteins (RNPs). WT RAW264.7 cells were subjected to two rounds of nucleofection with Cas9-RNPs using the SF Cell Line 4D-Nucleofector X Kit S (Lonza) and the 4D Nucleofector apparatus precisely as specified by the manufacturer. *Ifnar1* and *Ifnar2* gRNAs were delivered in the first and second rounds of nucleofection, respectively; both *Mavs* gRNAs were delivered in first and second rounds. *Ifnar* deletion was confirmed by luciferase assay as described under “Luciferase assay” in response to 8 hrs stimulation with titrated recombinant mammalian IFNB (PBL, 12405); *Mavs* deletion was confirmed by western blot.

### Suppression of LINE1 using CRISPR-dCas9-KRAB

In brief, two lentiviral vectros were used to target endogenous LINE1 elements in RAW264.7 macrophages. gRNAs were cloned into the puro-resistant pKLV2-U6gRNA5(BbsI)-PGKpuro2ABFP-W(Tzelepis et al., 2016) vector. A second vector expressing dCas9-KRAB under control of the SFFV promoter was co-transduced into cells(Gilbert et al., 2014) for epigenetic suppression of gRNA targets. gRNAs against LINE1 5’ regions were generated using CRISPOR and selected based on their predicted *in silico* enrichment in full length LINE1 elements and paucity of genic targets.

### Generation of human monocyte derived DCs (MDDCs)

Buffy coats were prepared from peripheral adult human blood and CD14^+^ monocytes were isolated by magnetic separation (Miltenyi 130-050-201). CD14^+^ monocytes were differentiated into DCs (MDDCs) in RPMI containing Glutamax, supplemented with 10%FBS (GIBCO), 50µg/ml Gentamicin (GIBCO), 0.01M HEPES (GIBCO), 10ng/ml GM-CSF (Miltenyi premium grade), and 50ng/ml IL-4 (Miltenyi premium grade) at a density of 10^6^ cells/ml. Cells were harvested on day 4 for further analyses.

### Lentiviral transduction of human cells

Lentiviral particles were produced as previously described(Gentili et al., 2019). Briefly, 293FT in one well of a 6-well were transfected with 1µg psPAX2, 0.4µg pCMV-VSV-G and 1.6µg of a lentiviral vector (for human shSUV39H1, Sigma, TRCN0000157251, TRCN0000275323, TRCN0000275322) using TransIT 293 (Mirus, Biomedex) in Optimem (GIBCO). 2.6µg of pSIV3 and 0.4µg of pCMV-VSV-G were transfected into 293FT cells to generate SIV-VLPs. Twelve to fourteen hours later, the medium was changed to 3 mls of MDDC culture medium without cytokines. 30-32 hours later, supernatants were harvested, passaged over a 0.45µm filter, and used immediately for transduction of freshly isolated CD14^+^ monocytes at 1.5x10^6^ cells/well of a 6-well plate in 1ml of MDDC culture medium. 1 ml of each lentiviral particles and SIV-VLPS were added to each well for 3mls final volume in the presence of protamine (8µg/ml). 48hrs after transduction, puromycin (2µg/ml, Invivogen) was added for seletion of shRNA-expressing cells. Cells were harvested on day 4 of culture, plated at 50,000 cells per well of a 96-well round bottom plate, and stimulated overnight with the indicated PAMPs.

### Luciferase assay

Production of type-I IFN production in RAW264.7 macrophages was measured by production of secreted luciferase under the control of the interferon-inducible ISG54 promoter. Briefly, conditioned supernatants were collected after stimulation, and 10µl was quantified for secreted luciferase (Renilla) activity in the presence of luciferase substrate (Quanti-Luc; rep-qlc2, Invivogen). Luminescence was recorded on a FLUOstar OPTIMA microplate reader (BMG Labtech).

### IFN ELISA

IFNb in conditioned supernatants was measured with the VeriKine Mouse Interferon Beta ELISA Kit (PBL, 42400-1) precisely as specified in the manufacturer’s protocol.

### RNA purification, Sequencing, and qPCR

Following stimulation, BMDCs were harvested on ice, collected by centrifugation and immediately dissolved in Trizol (Life Technologies, 15596018) and stored at -80°C until further processing. RNA was separated from DNA and protein by chloroform extraction. Briefly, 100µl chloroform was added to 1 ml of Trizol, mixed by shaking for 1 min and incubated for 5 minutes at RT. After centrifugation (20,000 x *g* for 25 minutes at 4°C), 400µl of the RNA-containing aqueous phase was removed and combined with 1µg of RNAse-free glycogen (ThermoFischer) and 500µg ice-cold isopropanol. The mixture was frozen overnight at -80°C and thawed on ice before centrifugation (20,000 x *g* for 25 minutes at 4°C). The RNA pellet was air-dried for 30-60minutes in a laminar hood, resuspended in 10µl RNAse/DNAse-free water, and the DNA removed by in-solution DNAse digestion (Qiagen’s RNase-Free DNase Set, #Cat 79254) for 30 minutes at room-temperature. The resulting DNA-free RNA prep was further purified over Quiagen columns using the Qiagen RNeasy mini kit and eluted in RNase/DNase-free water. RNA preparations performed with this protocol routinely yielded an A260/280 ratio above 2 and averaging 2.08. HTS libraries were constructed using the TruSeq Stranded mRNA kit from Illumina and sequenced to a depth of at least 50x10^6^ reads per sample using 100bp paired-end sequencing.

For qPCR, cDNA was generated from 500ng of RNA using random hexamers (Promega, C1181) and the SuperScript III First-Strand kit (Themro Fisher, 18080051) in a 20µl reaction. 2µl of RNaseH was added and the reaction carried out at 37°C for 20 minutes. Following cDNA synthesis, 160µl of RNAse/DNase-free H_2_0 was added, and 4µl of this was used per qPCR reaction containing 1µl of 5µM primers and 5µl of SybrGreen reagent (LifeTechnologies, 4367659). 10µl reactions were performed in a 384 well plate (Thermo Fisher, 4483319) on a Viia7 thermal cycling system (Applied Biosystem).

### PolyA^+^ RNA-Seq Analysis Sequencing Data Processing

FASTQC [http://www.bioinformatics.babraham.ac.uk/projects/fastqc/.] was run on all samples to assess the quality of sequencing. Raw sequences were mapped to the mouse mm10 reference genome (Refseq) with STAR aligner (Spliced Transcripts Alignment to a Reference, version 2.5.0a) (Dobin et al., 2013). We used the option *--bamRemoveDuplicatesType UniqueIdentical* to mark all multimappers, and duplicate unique mappers, *--outFilterMultimapNmax 100* that correspond to the number of read alignments which will be output only if the read maps fewer than this value.

### TE Quantification and Intactness

We used the Homer Software as previously described. However, we considered only TEs for the analysis if the row sum across all samples was FPKM 5. LINE1 and LTR elements greater than or equal to 4kb in length were considered intact. SINE elements were considered intact when they diverged less than 10% from their consensus sequences.

### Differential Analysis

Using the raw reads matrix, we filtered out the non-expressed genes and TE separately by requiring more than 5 reads in at least 2 samples for each features and normalized using DESeq2 R-package v1.18.1 (Love et al., 2014). Then, the differential expression analysis was performed using DESeq2, only the Benjamini Hochberg (BH) adjusted p-value below 0.05 were considered as significant.

### PCA

The PCA was performed using the Ade4 package(S Dray, 2007) of the R software (version 3.3.2). The barycenters were computed from the set of observations in each condition and projected into the PCA plot.

### Chromatin Immunoprecipitation (ChIP)-sequencing

Histone ChIP experiments were performed as previously specified (Blecher-Gonen, et al 2013) with several changes. Briefly, harvested BMDCs were collected by centrifugation at 350 x *g* for 5 minutes and resuspended in 1ml of I-10 medium. Cells were cross-linked with 1% formaldehyde (Euromedex) for 8 minutes at room temperature and the reaction quenched in 150mM glycine for 10 minutes. Cross-linked cells were collected by centrifugation (350 x *g* for 5 minutes at room temperature) and washed twice with cold PBS. Pelleted cells were subsequently lysed to isolate cross-linked chromatin. For cytosolic extraction, cells were first suspended at 1x10^6^ cells/100µl of lysis buffer 1 (LB1) (50mM Hepes (pH7.5), 140mM NaCl, 1mM EDTA (pH 8.0), 10% glycerol, 0.75% NP-40, 0.25% Triton X-100) with protease inhibitors (Roche 11873580001) and Na_3_VO_4_ (180µg/ml) for 10 minutes on ice, followed by centrifugation at 500 x *g* for 5 minutes at 4°C to separate cytosol and nuclei. Nuclear pellets were then dissolved in lysis buffer X (LBX) (50mM Tris pH 8.0, 5mM EDTA pH 8.0, 0.25%SDS) supplemented inhibitors at 10^6^cells/100µl and incubated on ice for an additional 10 minutes, and then sonicated with a Bioruptor Pico (Diagenode) for 11 cycles (30 seconds on/off) at 4°C. Lysates were cleared by centrifugation at 20,000 *x g* for 20 minutes at 4°C, snap frozen on liquid N_2_, and stored at -80°C until immunoprecipitation. Samples were thawed on ice and diluted 1:1 with dilution buffer X (DBX) (50mM Tris pH8.0, 0.9%NP40, 350mM NaCl), followed by the addition of either anti-H3K9me3 (Diagenode) or control rabbit IgG (Diagenode) and rotated overnight at 4°C. The following day, 20µl of DBX-washed Magna ChIP Protein W+G magnetic beads (Millipore) were added to each sample and rotated for 1hr at 4°C, followed by washes and de-crosslinking exactly as described previously(Blecher-Gonen et al., 2013). DNA fragments were purified by two-sided selection with SPRI beads (Beckman Coulter B23318) according to the manufacturer’s guidelines and Illumina sequencing libraries prepared with the TruSeq ChIP Library Preparation Kit. 150bp paired-end sequencing of pooled libraries was performed on a HiSeq 2500 (Illumina) platform in rapid run mode to yield 50x10^6^ reads per sample.

### Cross-linked cytosolic and nuclear cGAS immunoprecipitation and next generation sequencing (NGS)

Cytosolic cGAS immunoprecipitation was carried out similar to histone ChIP experiments with minor adjustments. Anti-GFP (ChromoTek) or control beads (ChromoTek) were added to the appropriate cell number as indicated by the manufacturer and incubated overnight with rotation at 4°C. The following day, samples were collected with a magnet, transferred to a 96-well qPCR plate and washed, de-crosslinked and purified as for histone ChIP. Purified cGAS-bound DNA was sheared to a mean size of 200bp in 100µl of direct elution buffer (10 mM Tris-HCl (pH 8.0), 5 mM EDTA (pH 8.0), 300 mM NaCl and 0.5% SDS (vol/vol) with a Covaris S220. Sheared DNA was purified with Agencourt AMPure XP beads (Beckman Coulter). Illumina-based libraries were prepared to preserve ssDNA, RNA:DNA hybrids, and dsDNA. Briefly, DNA fragments were tailed and ligated with an adaptor, followed by primer-based PCR extension of complementary DNA. A second adaptor was ligated to the 5’ end of the template strand. dsDNA species were amplified and indexed using barcoded primers for Illumina-based high throughput sequencing, and 150bp paired-end sequencing of pooled libraries was performed on a HiSeq 2500 (Illumina) platform in rapid run mode to yield 50x10^6^ reads per sample.

### Mapping

ChIP-seq reads were mapped to mouse genome build mm10 with Bowtie2 (v2.1.0) (Langmead and Salzberg, 2012)using the global alignment mode allowing 1 mismatch in seed alignment. We processed only mapped reads.

### Transposable elements and genes quantification

For computing quantification of TE and gene levels, genome-mapped reads were used with Homer Software (using analyzeRepeats.pl script)(Heinz et al., 2010). We used different options : (i) –*noCondensing* to report the number of reads for each repeat element separately or (ii) –*condenseL1* to condense reads from each repeat name on one entry. Finally, the number of reads mapped to each TE was normalized to its length and total number of genome-aligned reads (FPKM value, Fragments Per Kilobase of exon model per Million mapped reads using (-rpkm). TE and genes with RPKM0.1 were excluded. Using these parameters, 85,795 unique TEs and genes were kept for analysis.

### Human MDDC RNAseq dataset

Publicly available RNA seq datasets from untreated and LPS stimulated human MDDCs were downloaded from ENCODE (GSE94180) and aligned to the human genome using the GRCh38/hg38 assembly.

### ChIP-Seq Analyses Mapping

H3K9me3 ChIP-seq reads were mapped to mouse genome build mm10 with Bowtie2 (v2.1.0) using the global alignment mode allowing 1 mismatch in seed alignment (Langmead and Salzberg, 2012). We processed only mapped *reads* with high mapping quality (*MAPQ>=20*) using samtools (v0.1.8).

### Transposable elements and genes quantification

For computing quantification of TEs and genes levels, we used genome-mapped reads were used with Homer Software (v4.7) (using analyzeRepeats.pl script) (Heinz et al., 2010). We used different options : (i) –noCondensing to report the number of reads for each repeat element separately or (ii) -condenseL1 to condense reads from each repeat name on one entry. Finally, the number of reads mapped to each TE was normalized to its length and total number of genome-aligned reads (FPKM value, Fragments Per Kilobase of exon model per Million mapped reads using (-rpkm).

### Peak Calling and Annotation

Peak calling was performed with SICER v1.1 using parameters : ’-W 200 -G 600’, peaks at 1 % FDR threshold(Xu et al., 2014). Annotation of Peaks was performed using Homer Software (with AnnotatedPeaks.pl script).

### Bigwig files and coverage visualization

All Bigwig files were generated with merged biological replicates to improve the sensitivity by increasing the depth of read coverage and normalized over own input ChIP-seq files using -- BamCompare (with options: --effectiveGenomeSize 2719482043 for mm10 and --binSize 10 and –normalizeUsing BPM) from deepTools (v3.1.1) (Ramirez et al., 2014). For heatmap visualization were generated with --PlotHeatmap after to compute the matrix of scores per genome regions generated by the tool --ComputeMatrix from deepTools Software. Steady state H3K9me3 Suv39h1-dependent elements in WT vs KO (Figure 1C) were selected using *bedtools intersect* (default parameters) with WT peak and *subtract* (-A) with KO peak. H3K9m*e3 Suv39h1*-dependent elements in untreated vs LPS-treated (2hr and 4hr) were selected using *bedtools intersect* (default parameters) with UT peak and *subtract* (-A) with WT 2hr peak and WT 4hr peak.

### PAGE and Western Blot

For some proteins including SUV39H1, cells were lysed with LBX supplemented with EDTA-free protease inhibitors (Roche), phosphatase inhibitors (Halt phosphatase, ThermoFischer), and benzonase. They were incubated on ice for 30 minutes, vortexed at high speed for 15 seconds and then centrifuged for 10 minutes at 13,000x*g,* aliquoted, snap frozen on liquid nitrogen and stored at -80°C until further processing. Samples were thawed on ice and denatured in Laemmli sample buffer supplemented with fresh beta-mercaptoethanol for 15 minutes at 95°C. For phosphorylated and corresponding total protein levels including IRF3 and STING, cells were lysed in ice cold RIPA buffer (50mM Tris HCL, 150mM NaCl, 0.1% SDS, 0.5% DOC, 1%NP-40, EDTA-free protease inhibitor (Roche), phosphatase inhibitors (Halt phosphatase, ThermoFischer), and benzonase (ThermoFisher)). Lysates were incubated for 30 minutes on ice, vortexed for 15 seconds at high speed, and centrifuged for 8 minutes at 8,000 x *g*. Clarified lystates were aliquoted, snap frozen on liquid nitrogen and stored at -80°C. Samples were thawed on ice and denatured in Laemmli sample buffer supplemented with fresh beta-mercaptoethanol for 15 minutes at 95°C. Lysate from 10^5^ cells were separated on 4–15% Mini Protean TGX Stain-Free gels (BioRad) and dry transferred to PVDF membranes (Bio-Rad) with the Trans-Blot Turbo Transfer System (Bio-Rad). In experiments where phosphor-STING was measured, lysate from 8.5x10^5^ cells was loaded per lane. Blots were blocked with TBS 0.05% Tween20 (TBST) and 5% non-fat milk (Carnation, total protein) or 10% Roche blocking reagent (phosphor-proteins) for 1hr at RT, rinsed in TBST, and then rocked overnight at 4°C in TBST 5% BSA (Fraction V, 04-100-812-C, Euromedex) with primary antibodies: cGAS (1:1000; 31659, CST), gp96 (1:1000; adi-spa-850-D, Enzo), Phosphor-IRF3 (Ser396)(1:1000; 29047, CST), Phosphor-STING (1:1000; CST), Phosphor-NF-KappaB p65 (Ser536)(1:1000; 3033, CST), Phosphor-TBK1/NAK (Ser172) (1:1000; 5483, CST), SUV39H1(1:500; 8729, CST), TBK1 (1:1000; 3504, CST), anti-RAB6 (1:1000, sc-310; Santa Cruz Biotechnology), Histone H3 (1:1000, CST), MDA5 (1:1000, CST). The following morning, blots were washed three times for ten minutes each in TBST at RT and then rocked for 1hr at RT in blocking buffer containing secondary antibodies (1:10,000; Jackson ImmunoResearch): anti-rat-horseradish-peroxidase (HRP) (112-035-143) or anti-rabbit-HRP (111-036-046). Following 4 more washes, membranes were coated with an HRP substrate (Enhanced Chemiluminescence, 32106; Thermo Fisher Scientific) and the images captured using a BioRad Chemidoc imager.

### Flow Cytometry

Surface staining, flow cytometry, and washes were performed in FACS buffer (2%FBS, 0.5mM EDTA, PBS). BMDCs were surface-labeled with fluorochrome-coupled antibodies to the following antigens (all 1:400): MHCII-v450 (Invivogen, 48-5321-82), CD11b-PerCP-Cy5.5 (Invivogen, 45-0112-82), CD11c-PE-Cy7(Invivogen, 25-0114-82), CD86-APC(BD Biosciences, 561964). Dead cells were identified by APC-Cy7 live-dead (eBioscience, L34975) staining. For intracellular staining, BMDCs were fixed and permeabilized using the eBioscience FoxP3 staining kit according to the manufacturer’s recommendations, followed by incubation with anti-Viperin-PE (BD Biosciences,565196) with 1%normal mouse/rat serum (Sigma), and FC block (1:200; BD Biosciences, 553141) in permeabilization buffer for 45 minutes at RT. Cells were washed twice in permeabilizatoin buffer and a second time in FACS buffer before flow cytometric analysis. Flow cytometric data were collected on either a MacsQuant (Miltenyi) or a FACSVerse(BDbiosciences).

### Cytosolic DNA isolation and RTI

Cytoslic DNA from 2x10^6^ BMDCs was isolated as previously described (*42*). After LPS stimulation, BMDCs were harvested from 6-well dishes, washed twice in PBS, and resuspended in 200µl of digitonin buffer (150mM NaC, 50mM HEPES pH7.4, 25µg ml^-1^ digitonin (Sigma)). Samples were rotated end-over-end for 20 minutes at 4°C, cleared by centrifugation at 18,000 x *g* for 10 minutes to collect nuclei and cellular debris which were then dissolved in LBX and saved as the pellet fraction for normalization. Soluble material was treated with DNase-Free RNaseA (20µg) (Thermo Fisher, ENO531) at 37°C for 1hr, followed by proteinase K (20µg) (Thermo Fisher AM2548) for 1hr at 56°C. DNA was further purified with MinElute columns (Quiagen, 28004) or with AmpureXP beads (Beckman Coulter) according the manufacturer’s guidelines and eluted in 250µl of DNase/RNase-free H_2_O. 4µl of DNA extract from cytosol or the pellet fraction were amplified as described for qPCR of cDNA. For reverse transcriptase inhibitor treatment, a combination of emtricitabine, tenofovir disoproxil fumarate, and nevirapine were maintained at 10µM each in BMDC cultures from day 1 until the end of treatment. For control, DMSO vehicle was maintained alongside RTI-treated cultures.

### Lentivirus and shRNA

Lentiviral shRNA (pLKO.1) plasmids for mouse Suv39h1 (7µg) (Sigma TRCN0000097439, TRCN0000097440, TRCN0000097441, TRCN0000097443) or control shRNAs for GFP (Sigma) and luciferase (Sigma) were combined with 7µg of psPAX2 packaging and 0.7µg of CMV-VSVg envelope vectors in 1ml of OptiMEM medium. 45µl of room temperature Trans-IT 293 Transfection Reagent (Euromedex, Mirus 2700) was then added and the mixture incubated at RT for 30 minutes. All contents were then delivered to HEK293FT cells that were plated in 10mls of DMEM (Gibco), 10%FBS (VWR, 11543387), 1%Penicillin, Streptomycin (Gibco) at 2-3x10^6^ cell/10cm plate the previous night. The following morning, BSA (Sigma A7979) was added to a final concentration of 1% and the viral supernatants harvested 24hrs later by passaging through a 0.45µm filter. 2.5 mls of supernatant were then added to 0.5x10^6^ cells in mls in a 6-well tissue culture-treated dish () and left overnight. Cells were washed the next day and subjected to puromycin selection (2µg/ml, Invivogen ant-pr-5) 48hrs later. Knockdown efficiency was assessed by western blotting or qPCR as specified.

### SUV39H1 cloning and overexpression

For mouse SUV39H1 overexpression studies, a previously described lentiviral expression plasmid carrying RFP reporter and puromycin resistance genes (pL-SFFV.Reporter.RFP657.PAC, Addgene#61395) was used with the following modifications. After AgeI digestion, PacI and AbsI sites were introduced and one of the AgeI sites preserved downstream of the SFFV promoter by polylinker ligation using the following oligos: ACCGGTTTGGGATTAATTAAAATCACCTCGAGGCAGTCCGGT and TGGCCAAACCCTAATTAATTTTAGTGGAGCTCCGTCAGGCCA. In brief, oligos were phosphorylated with T4 PNK (NEB) for 30 minutes at 37°C, denatured at 95°C for 5 minutes and then ramped down to 25°C at 5°C per minute. Ligation proceeded overnight at 16°C using a thermocycler, and bacterial transformation of NEB 10-beta competent E. coli (NEB) accomplished by heat shock at 42°C. Minipreps were confirmed by Sanger sequencing.

Sequential digestion of modified pL-SFFV-RFP with AgeI and BspEI were carried out at 37°C with column purification between reactions (QIAquick, Quiagen). A mouse SUV39H1 gBlock was synthesized (IDT) and cloned into AgeI/BSpEI-digested pL-SFFV-RFP using Gibson assembly (NEB) for 1hour at 50°C. NEB10 (NEB) bacteria were transformed and selected on ampicillin plates after 14hrs at 37°C. Minipreps were grown at 30°C over night with shaking at 250rpm. Plasmids constructs were confirmed by Sanger sequencing. The final SUV39H1 plasmid contained a P2A signal followed by the RFP reporter gene, and an internal ribosome entry site (IRES), followed by the puromycin resistance gene.

### cGAS reconstitution

Human cGAS WT open reading frame was amplified by PCR from cDNA prepared from monocyte-derived dendritic cells and was previously described (Gentili et al., Science 2015). Murine cGAS WT open reading frame was amplified by PCR from cDNA prepared from C57BL6 murine bone-marrow derived dendritic cells and was previously described (Gentili et al., Science 2015). Human cGAS 161-522 C395A/C396A, and E225A/D227A were obtained by overlapping PCR. pTRIP-SFFV was generated by substitution of the CMV promoter with the SFFV promoter from GAE-SFFV-GFPWPRE and was previously described (Gentili et al., Science 2015, Raab et al., Science, 2016). Murine cGAS cDNA , Human cGAS 161-522 E225A/D227A, and Human cGAS 161-522 C395A/C396A were cloned in frame to EGFP in pTRIP-SFFV-EGFP to obtain pTRIP-SFFV-EGFP-ms cGAS, pTRIP-SFFV-EGFP-cGAS 161-522 C395A/C396A, and pTRIP-SFFV-EGFP-FLAG-cGAS 161-522 E225A/D227A. Leniviral particles were generated as described and used to transduce WT RAW264.7 macrophages.

### Software

Sequencing data was analyzed with R packages and R-Studio; heatmaps were generated using Morpheus software from the Broad Institute (https://software.broadinstitute.org/morpheus/). Flow cytometry data was analyzed with FloJo (Tree Star) v9.9.5. Graphs and statistical analysis were generated with Prism/Graphad (Version 7). Figure layouts were constructed with Prism/Graphpad, Adobe Illustrator, and Adobe Photoshop.

## Notes

### Competing Interest Statement

The authors have declared no competing interest.

